# Distinct specific interactions of the UapA transporter with membrane lipids are critical for dimerization, ER-exit and function

**DOI:** 10.1101/710897

**Authors:** Anezia Kourkoulou, Pothos Grevias, George Lambrinidis, Euan Pyle, Mariangela Dionysopoulou, Argyris Politis, Emmanuel Mikros, Bernadette Byrne, George Diallinas

## Abstract

Transporters are transmembrane proteins that mediate the selective translocation of solutes across biological membranes. Recently, we have shown that specific interactions with plasma membrane phospholipids are essential for formation and/or stability of functional dimers of the purine transporter, UapA, a prototypic eukaryotic member of the ubiquitous NAT family. Here, we show that distinct interactions of UapA with specific or annular lipids are essential for *ab initio* formation of functional dimers in the ER or ER-exit and further subcellular trafficking. Through genetic screens we identify mutations that restore defects in dimer formation and/or trafficking. Suppressors of defective dimerization restore *ab initio* formation of UapA dimers in the ER. Most of these suppressors are located in the movable core domain, but also in the core-dimerization interface and in residues of the dimerization domain exposed to lipids. Molecular Dynamics suggest the majority of suppressors stabilize interhelical interactions in the core domain and thus assist the formation of functional UapA dimers. Among suppressors restoring dimerization, a specific mutation, T401P, was also isolated independently as a suppressor restoring trafficking, suggesting that stabilization of the core domain restores function by sustaining structural defects caused by abolishment of essential interactions with specific or annular lipids. Importantly, introduction of mutations topologically equivalent to T401P into a rat homologue of UapA, namely rSNBT1, permitted the functional expression of a mammalian NAT in *A. nidulans*. Thus, our results provide a potential route for the functional expression and manipulation of mammalian transporters in the model Aspergillus system.

**Author Summary:** Transporters are proteins found in biological membranes, where they are involved in the selective movement of nutrients, ions, drugs and other small molecules across membranes. Consequently, their function is essential for cell viability, while their malfunction often results to disease. Recent findings have suggested that transporter functioning depends on proper interactions with associated membrane lipids. In this article, using UapA, a very well-studied transporter from a model fungus (*Aspergillus nidulans*), we show that two types of specific interactions with lipids are essential for tight and specific association of two UapA molecules in a single functional unit (UapA dimer), and for targeting to the cell membrane and transport activity. The first type of interaction concerns specific lipids associating with positively charged amino acids at the interface of the UapA dimer, whereas the other type involves lipids that interact with charged amino acids at the outer shell of the transporter. Most interestingly, defects due to abolishment of UapA-lipid interactions were shown to be restored by mutations that increase UapA stability. Using this information, we genetically manipulated and increased the stability of a mammalian transporter (rSNBT1), and thus achieved its functional expression in the experimentally tractable system of *A. nidulans*.

## Introduction

Transporters are essential transmembrane proteins that catalyze the uptake or efflux of metabolites, nutrients, ions and drugs across biological membranes. Transporter malfunction, due to genetic mutations or metabolic defects, results in significant cellular or organismal disruption [1] (http://www.tcdb.org/). Despite their biological and apparent medical importance, knowledge on structure-function relationships in transporters is limited compared to extramembrane hydrophilic proteins. This is in part due to complexity associated with their translocation and co-translational folding into a membrane lipid bilayer (the ER in eukaryotes or the plasma membrane in prokaryotes). Additionally, in eukaryotes, transporters follow specific membrane trafficking, turnover or recycling routes, which add further complications in understanding their mechanisms regulation of expression, function and turnover [2–6]. A further contributory factor is their functional and structural dependence on specific membrane lipids, something which is only recently being explored in detail [7, 8]. Further complications for transporter study arise from difficulties in expressing sufficient quantities for downstream structural studies, functional reconstitution in proteoliposomes, or in measuring their kinetics in intact cells where the presence similar transporters with overlapping specificities complicate the analysis [9].

One of the best studied eukaryotic transporters is the UapA xanthine-uric acid/H^+^ symporter of the filamentous ascomycete *Aspergillus nidulans* [10, 11]. This is due to development of rigorous genetic, biochemical and *in vivo* cellular approaches, uniquely available in the model system of *A. nidulans*, and more recently, by structural and biophysical studies. The high-resolution, inward-facing, structure of a conformationally locked mutant of UapA (G411V_Δ1-11_), together with genetic and other functional studies, revealed that UapA functions as a homodimer [12]. In the UapA dimer each monomeric unit consists of a core domain that hosts the substrate binding site and a dimerization domain that includes elements crucial for substrate specificity. Molecular Dynamics (MD) simulations together with comparison with other related proteins suggested that UapA functions via an elevator mechanism, as reported for a range of other transporters [13–18]. Upon substrate binding, the core domain moves against the relatively immobile dimerization domain to transport the substrate from one side of the membrane to the other [12, 19]. Interestingly, UapA substrate specificity is also determined by the proper formation of the dimer, as a specific Arg residue (Arg481) from one monomeric unit dynamically controls the substrate translocation trajectory in the opposite monomer [12]. Recently, native Mass Spectrometry (MS) of purified UapA, combined with MD, mutagenesis, and functional analyses, established that the membrane lipids, phosphatidylinositol (PI) and phosphatidylethanolamine (PE), have a critical role in stabilizing the functional UapA dimer [20]. More specifically, it has been shown that UapA delipidation during purification causes dissociation of the dimer into monomers, but subsequent addition of PI or PE both recovers lipid binding and re-forms the UapA dimer. MD simulations predicted possible lipid binding sites near the UapA dimer interface, and subsequent mutational studies confirmed that Arg287, Arg478 and Arg479 act as the lipid binding residues involved in the formation of UapA dimers and are absolutely necessary for transport activity [20]. Importantly, however, while triple-alanine replacement of these arginines leads to total lack of UapA function, both native MS and the Bifluorescence Complementation (BiFC) assays indicated that a fraction of UapA can still dimerize [20, 21]. Thus, lipid binding might not be an absolute requirement for dimer formation, but is essential for formation and/or stability of *functional* dimers. The total lack of function of the lipid binding site UapA mutant (R287A/R478A/R489A) further suggests that specific interactions with lipids are also necessary for the mechanism of transport *per se*. Rather surprisingly, the GFP tagged inactive triple arginine mutant can still properly traffic to the PM, which means that either monomeric UapA translocates normally to the PM, or that UapA dimer is initially formed in the ER and traffics to the PM, but then becomes unstable and dissociates into non-functional monomers. As MD simulations further predicted that lipids could also bind to the outermost, membrane-facing regions of the core domains of the UapA dimer [20] further investigation was needed to fully understand the role of membrane lipids in UapA folding, subcellular traffic and transport function.

Here we investigate further the role of residues Arg287, Arg478 and Arg479 in UapA dimer formation and/or stability, and study the role of additional interactions of UapA with annular lipids. Using mutational analyses and genetic screens for suppressors of mutations affecting putative lipid-binding residues, we confirm that that Arg287, Arg478 and Arg479 are essential for *ab initio* dimerization in the ER, but redundant for membrane traffic, whereas interactions between distinct residues (Lys73, Arg133 and Arg421) and annular lipids are essential for proper folding, ER-exit and membrane traffic. Our results reveal that genetic modification of residues in the core domain of UapA can compensate for the „lost‟ lipid interactions in the original mutants. Importantly, using information on a specific core residue that proved to have a key role in stabilizing UapA, Thr401, we genetically manipulate and achieve in functionally expressing, for the first time, a mammalian homologue of UapA in *A. nidulans.* This opens the way for functional expression of mammalian NAT transporters, including those essential for vitamin C transport in humans [22.23], in the model *Aspergillus* system.

## Results

### Residues Arg287, Arg 479 and Arg479 are crucial for *ab initio* dimerization of UapA in the ER

We have previously shown that arginine residues 287, 478 and 479 are essential for phospholipid-dependent functional dimerization of UapA at the PM [20]. To further understand the basis of the functional defect in the triple R287A/R478A/R479A mutant, here we examined whether loss of dimerization occurs *ab initio* at the level of the ER, or whether what we have previously observed was due to instability of UapA dimers at the PM. For this, we used a previously described BiFC assay [21], adapted to follow the sorting and subcellular localization of *de novo* made UapA. This is based on time-course experiments following the subcellular localization of *de novo* made *alcAp*-UapA-GFP, which showed that after 1 h of transcriptional derepression UapA-GFP is hardly visible, but at 2-3 h it labels the ER and at 4 h appears mostly in the PM [6]. Using this system, we followed reconstitution of split-YFP, via UapA dimerization, in a strain containing two copies of the *alcAp-uapA* gene, tagged either the N- or the C-part of the *yfp* orf [21]. To follow *de novo* made UapA in young mycelia, we repressed the transcription of *alcA_p_*-*uapA-yfp_n_* and *alcAp*-*uapA-yfp_c_* overnight (16 h in MM at 25 ° C in glucose MM), a period to allow conidiospores germination and young hyphae development, and then shifted the culture to fructose-derepression medium for 1-4 h of growth. We performed this assay using wild-type UapA and the triple R287A/R478A/R479A mutant. Fig 1A shows that in the wild-type control strain our assay detects early reconstitution of split-YFP fluorescence at the ER network at 3h, but fails to do so in the triple R287A/R478A/R479A mutant, where only a weak signal is observed. After 4h expression, the totality of wild-type UapA fluorescent signal marks the PM, whereas a weak cytoplasmic fluorescent signal and very low cortical localization is observed in the R287A/R478A/R479A mutant. This shows that Ala substitutions of the three Arg led to significant reduction of apparent UapA dimerization at the ER membrane, and further suggests that specific contacts with ER lipids might be a prerequisite for dimerization. Surprisingly, the interactions of R287, R478 and R479 with lipids and dimerization proved redundant for sorting of the mutant UapA to the PM, as judged by the normal PM localization of R287A/R478A/R479A tagged with intact GFP (Fig 1B). These findings suggest that in the R287A/R478A/R479A mutant non-functional UapA monomers or partially misfolded dimers can still be secreted to the PM.

**Fig 1.**
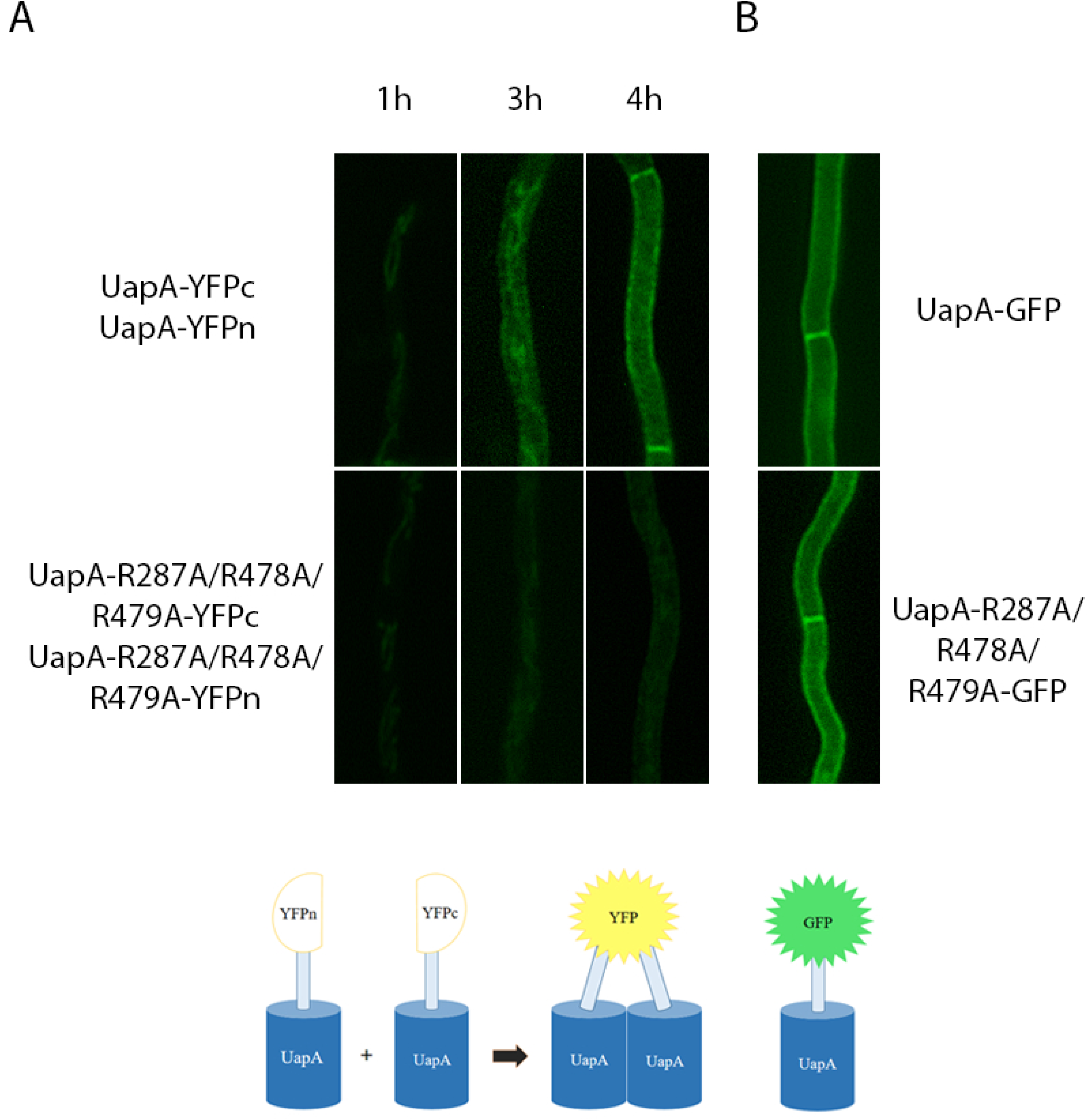
Residues Arg287, Arg 479 and Arg479 are crucial for *ab initio* dimerization of UapA in the ER. (A) Bimolecular complementation (BiFC) analysis based on reconstituion of split-YFP of *de novo* expressed wild-type or R287A/R478A/R479A mutant, expressed via the *alcA_p_* regulatable promoter in the presence of derepressive carbon and nitrogen two copies of the *uapA* gene, one tagged with the N-terminal and the other with the C-terminal part of the *yfp* orf (UapA-YFPc:UapA-YFPn and UapA-R287A/R478A/R479A-YFPc:UapA-R287A/R478A/R479A-YFPn). Notice the progressive appearance of a clear fluorescent signal in the wild-type UapA, firstly associated with ER membrane network (3h) and finally at the PM (4h). In contrast, in the R287A/R478A/R479A mutant, fluorescence remains extremely low, just above the level of detection. (B) Localization of wild-type or R287A/R478A/R479A mutant UapA tagged with GFP after 4h of transcription derepression, showing that the mutant can normally translocate in the PM despite very low apparent ability to dimerize, as shown in A. The lower panel shows schematically that dimerization UapA is required to detect a fluorescent signal from reconstituted split-YFP, while fluorescent signal from GFP-tagged UapA does not distinguish monomers form dimers.

### Genetic suppressors of R287A/R478A/R479A map in the core or dimerization domains

In order to further understand the molecular basis of how UapA-phospholipid interactions affect functional dimerization of UapA, we isolated genetic suppressors restoring UapA-mediated growth on uric acid in the R287A/R478A/R479A triple mutant, at 25 ° C, a temperature where this mutant does not grow on media containing UapA substrates. We purified 38 apparent suppressors and sequenced the orf. of the *uapA* gene. All 38 contained the original mutation (R287A/R478A/R479A) plus an extra point mutation, apparently the one that suppresses the growth defect on uric acid. S1 Table summarizes the identity and frequency of isolation of all suppressors, which concerned 13 distinct single amino acid substitutions in 11 different residues. Eight of the 13 suppressor were isolated more than once showing that mutagenesis was fairly saturated and also confirming that the amino acid changes detected are responsible for suppression. Fig 2A shows the topology of suppressor mutations in the UapA structure.

**Fig 2.**
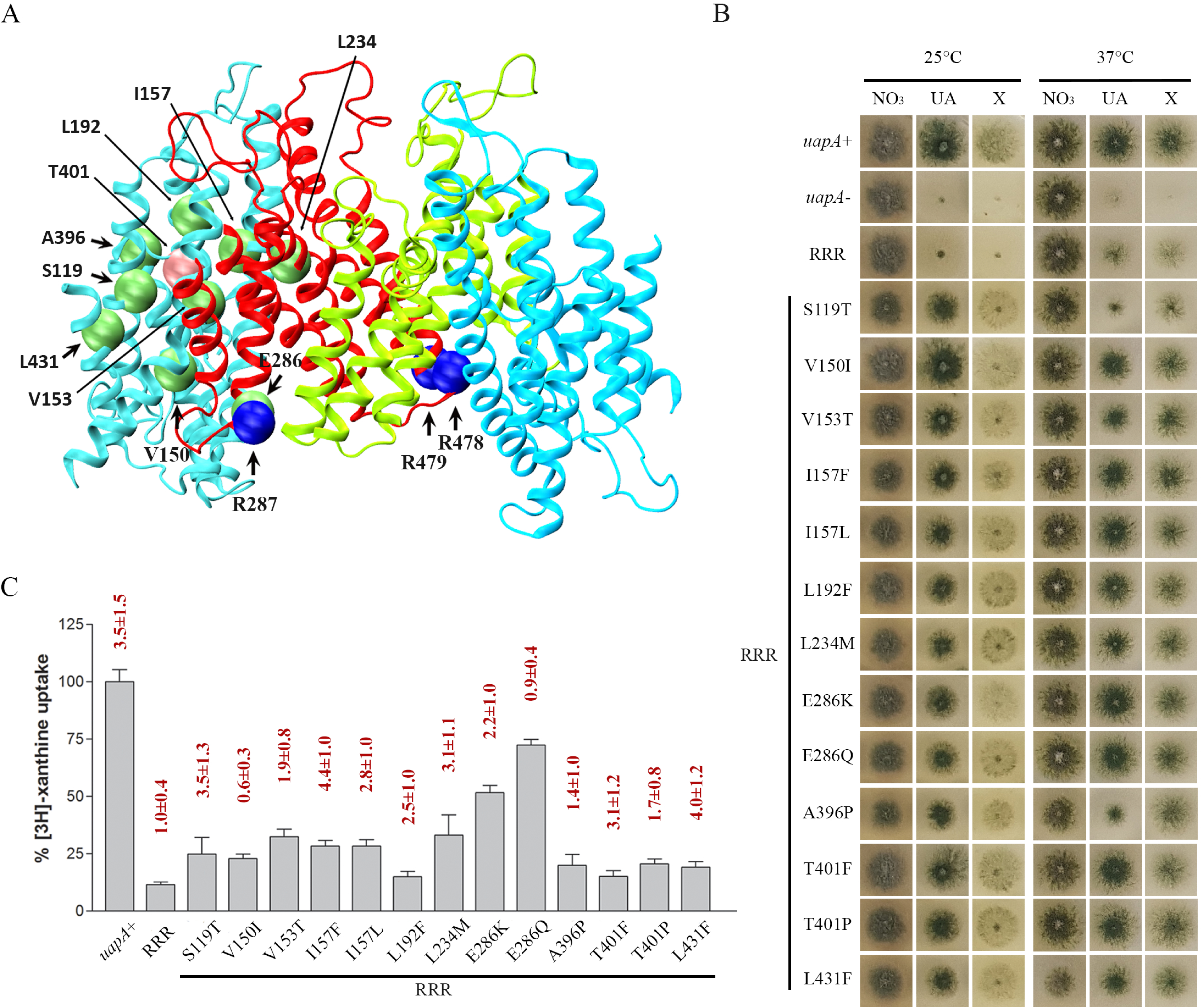
Genetic suppressors of R287A/R478A/R479A localized in the core and the dimerization domain of UapA partially restore UapA function. (A) Topology of amino acids modified in R287A/R478A/R479A suppressors. Core domains are colored light blue and dimerization domains red and green for clarity. Mutated amino acids in the original mutant strain are shown with blue spheres and in the suppressors with green and pink (T401) spheres. (B) Growth tests of R287A/R478A/R479A suppressors on UapA physiological substrates. Control strains are a strain with total genetic deletions in all major purine transporters (ΔACZ; negative control), referred in the figure as *uapA-*, and a ΔACZ transformant expressing wild-type *uapA–gfp* (*uapA+*; positive control). All suppressor strains and the original R287A/R478A/R479A strain are isogenic to the negative and positive control strains and express UapA from single-gene copies *uapA* tagged with gfp. All strains were grown in minimal media containing 10 mM nitrate (NO3), 0.5 mM uric acid (UA) or 1 mM xanthine (X) as N sources at 25°C (left panel) or 37°C (right panel). RRR depicts the R287A/R478A/R479A original genetic background. (C) Relative [^3^H]-xanthine transport rates of R287A/R478A/R479A and R287A/R478A/R479A suppressors expressed as percentage of initial uptake rates (V) compared to the wild-type (*uapA+*) rate. *K*_m_ values (μΜ) for xanthine are shown at the top of histograms. Results are averages of three measurements for each concentration point. SD was 20%.

The 13 distinct suppressors grew well, albeit slightly less than an isogenic strain expressing wild-type UapA, on uric acid or xanthine (Fig 2B). Among the suppressors, S119T scored as a thermosensitive mutant, growing very weakly on both UapA substrates at 37 ° C, similar to the original R287A/R478A/R479A. Direct uptake assays, measuring the transport rate of radiolabeled xanthine [24], were used to estimate the effect of the suppressor mutation on UapA transport kinetics. Fig 2C shows that in most suppressors UapA transport rates were re-established at ∼15-30% of the wild type protein, a level known to be the threshold for conferring visible UapA-mediated growth on uric acid or xanthine. Highest transport rates were obtained in suppressors E286Q (∼70%) and E286K (∼51%). In fact, the relative uptake differences of some suppressors (L192F, A396P, T401P, T401F or L431F) compared with the original R287A/R478A/R479A mutant were marginal. It should be noted that in growth tests purines are added to concentrations at the 1-2 mM level in order to be used as N sources, while in uptakes radiolabeled xanthine is used at sub-micromolar range (0.3-0.5 μΜ) for technical reasons. Thus any mutation that causes significant reduction in substrate affinity might score as an apparent loss-of-function mutation in uptakes, but still can allow normal growth on the relative substrate when this is supplied at mM concentration. To test whether the low apparent transport capacity of suppressors is due to reduced affinity for xanthine, we estimated the approximate *K*_m_ of several of the suppressors relative to the original R287A/R478A/R479A mutation or a wild-type UapA control. We found no significant reduction of affinities for xanthine in all suppressors tests (Fig 2C, on top of histograms).

### Genetic suppressors R287A/R478A/R479A re-establish UapA dimerization

As phospholipid binding has been shown to be essential for formation of functional UapA dimers, the suppressors of R287A/R478A/R479A could either restore functional dimerization, or confer transport activity to UapA monomers. To test these two alternatives, we performed Bifluorescence Competitions (BiFC) assays to follow UapA dimerization of *de novo* made UapA in selected suppressors (I157F, L234M, and T401P) and control strains, as described earlier for the original R287A/R478A/R479A mutant. Fig 3 shows significant reconstitution of split-YFP fluorescence, and thus apparent dimerization, at both ER and the PM, in all three suppressors studied, a picture similar to the wild-type. This confirms that suppressors restored function by restoring early dimerization of UapA at the ER.

**Fig 3.**
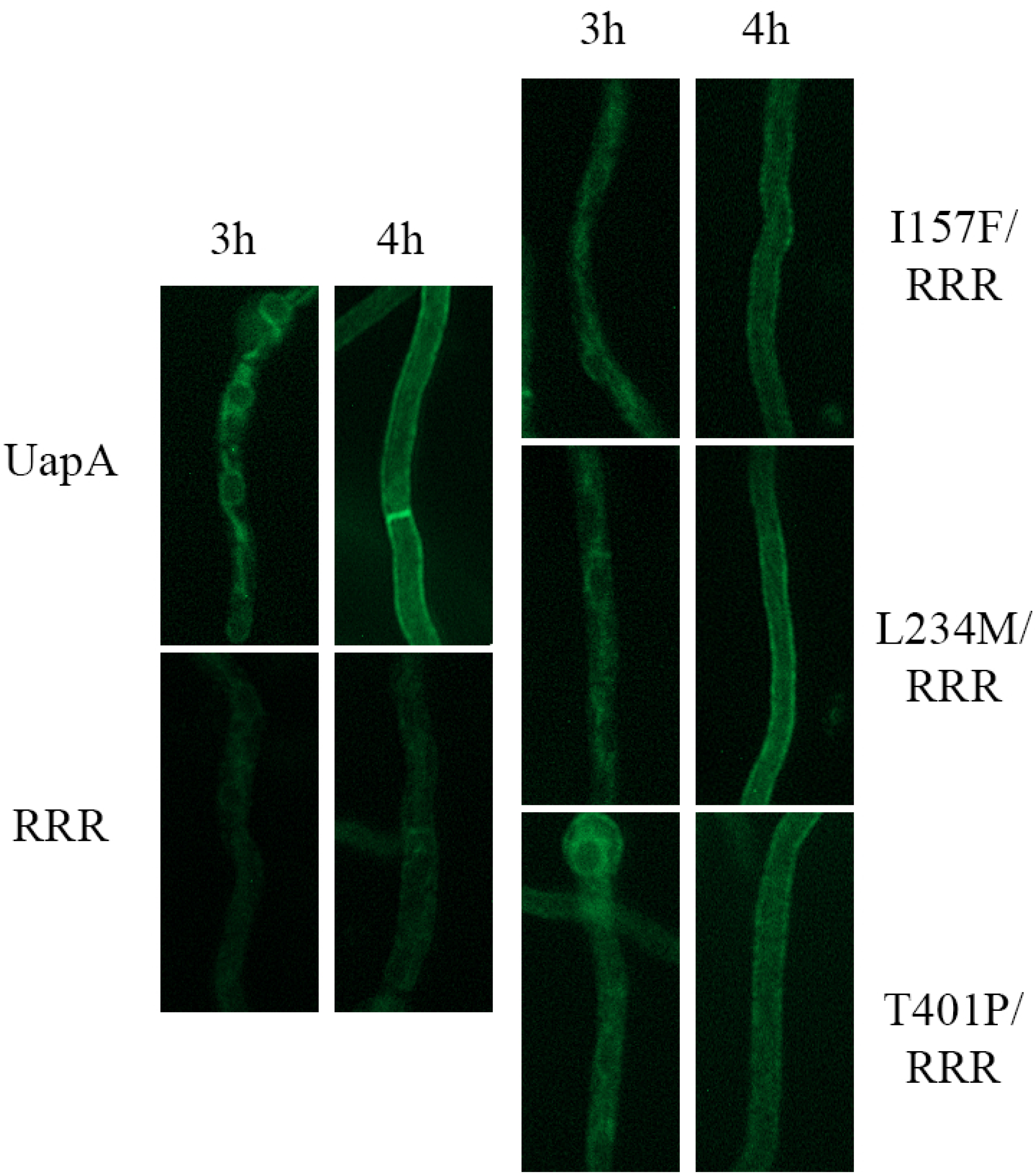
Genetic suppressors R287A/R478A/R479A re-establish UapA dimerization. Bimolecular complementation (BiFC) analysis based on reconstituion of split-YFP of *de novo* expressed UapA, R287A/R478A/R479A and selected R287A/R478A/R479A suppressors, namely I157F, L234M and T401P. Details of expression are as in Fig 1. Notice that all suppressors analyzed clearly restore, at least partially, the strength of the fluorescent signal detected in the ER and the PM, when compared to the original R287A/R478A/R479A mutant (RRR).

### Molecular Dynamics provide a structural rationale of the effect of suppressor mutations on UapA stability and function

Rather surprisingly, 10 out of the 13 mutations concerned residues located in transmembrane segments of the movable core domain (i.e. in TMS2, TMS3, TMS4, TMS9, TMS10 or TMS11). The core domain consists of two layers of transmembrane helices, the TMS1, TMS3, TMS8 and TMS10 in the inner part in contact with the dimerization domain, and TMS2, TMS4, TMS9 and TMS11 in the outer part, in contact with the membrane lipids. These two layers are stabilized through an extended network of hydrophobic interactions and certain key polar interactions which are mainly related with Asp388. Most suppressor mutations are related with the network of the hydrophobic interactions and are located on a virtual diagonal intersecting the core domain in the space between the two transmembrane layers (Fig 4 A, B). The UapA crystal structure (pdb 5i6c) shows that Leu192 interacts with Tyr189 (TMS4), Thr401 (TMS10), Ile157 (TMS3), Phe165 (TMS3), Ile101 (TMS1), Ile193 (TMS4), and Ile346 (TMS8). Similarly, Ile157 interacts with Ile101 (TMS1), Thr401 (TMS10), Pro402 (TMS10), Val349 (TMS8), Ile346 (TMS8) and Leu192 (TMS4). Thr401 is located in the center between the two TMS layers of the core domain and interacts with Tyr189 (TMS4), Leu192 (TMS4), Ile193 (TMS4), Val153 (TMS3), Ile157 (TMS3) and Ile101 (TMS1). Val153 is surrounded by Thr401 (TMS10), Pro97 (TMS1), Val94 (TMS1), Met400 (TMS10), Ser119 (TMS2), Met403 (TMS10) and Cys123 (TMS2). In the opposite side, Ser119 side chain is located in the middle between Val153 (TMS3), Val94 (TMS1), Met400 (TMS10) and Cys123 (TMS2). Finally, Leu431 is located between with Ala87 (TMS1), Met90 (TMS1) and Leu120 (TMS2). Contrary to the above mentioned residues, Leu234 is located in the dimerization domain, interacting mainly with Ile158 in TMS3. The suppressor mutations and more particularly L192F, T401F, L431F, S119T and V153T seem to enhance the above mentioned interactions and stabilize mainly the core domain. In order to verify this hypothesis, models of UapA including selected suppressor mutations were constructed and subjected to geometry optimization and short MD calculations. As shown in Fig 4C, D, E, F the phenyl moieties of the mutated residues I157F, T401F and L192F are accommodated in the space between the other lipophilic residues increasing hydrophobic interactions between TMS8, TMS3, TMS4 and TMS10. The introduction of a methyl group in the mutation S119T will also enhance, albeit to a lower degree, interactions with Val153 (TMS3) and Met400 (TMS10). Finally, the polar mutation V153T introduces a new hydrogen bond with Ser119 which already interacts with the backbone of Val94, creating a hydrogen bond network between TMS3, TMS2 and TMS1.

**Fig 4.**
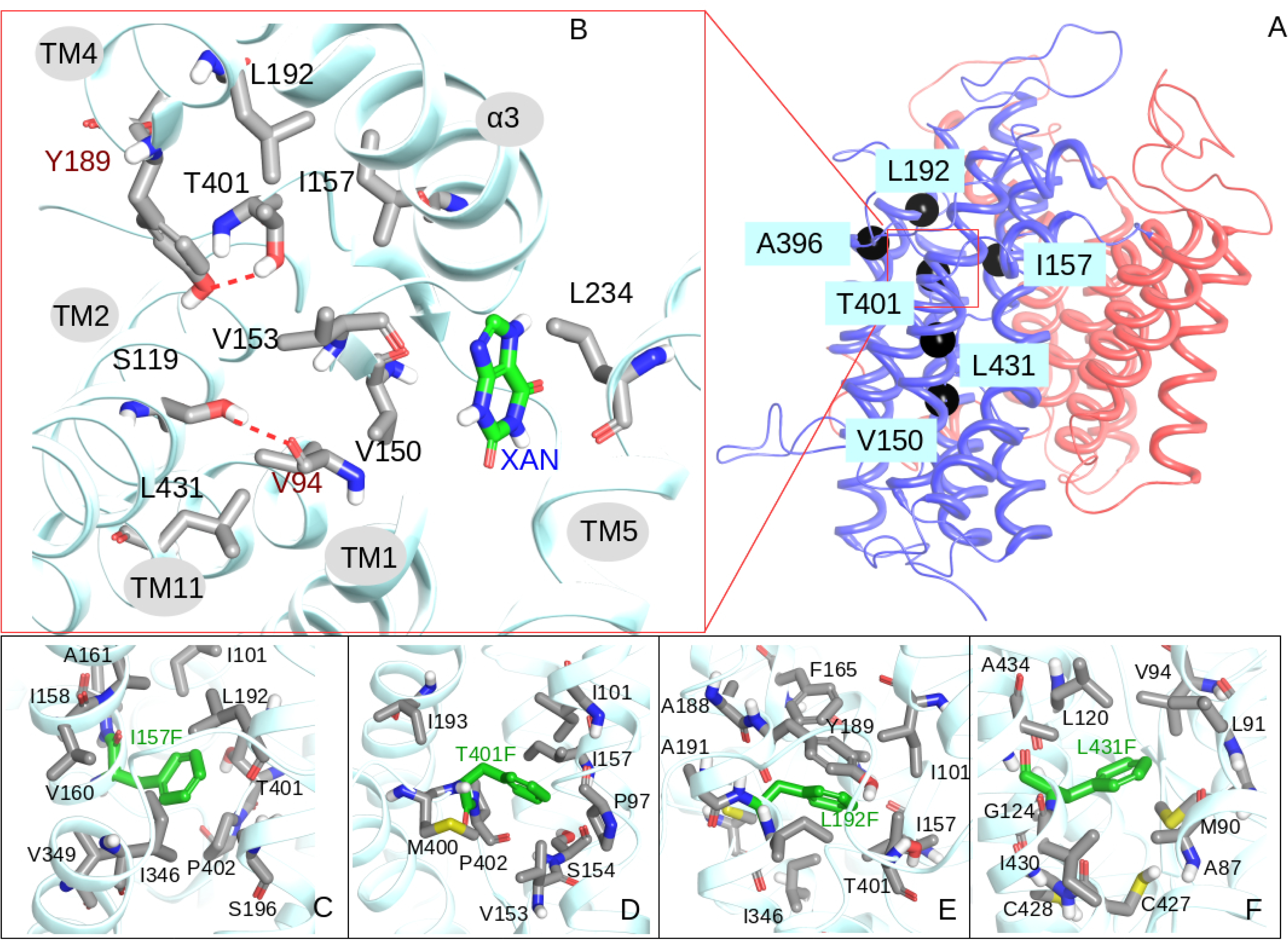
Molecular Dynamics provide a structural rationale of the effect of suppressor mutations on UapA stability and function. (A) Ribbon representation of UapA monomer. The core domain is colored blue and the gate domain red. Topology of Type I Suppressors on UapA crystal structure is depicted with black spheres. (B) Detailed view of the topology of T401 and amino acids within 4 Å, including residues mutated in suppressors in black lettering (see text). The substrate (xanthine) is also shown. (C-F). Detailed view of I157F, T401F, L192F and L431F mutation and amino acids within 4 Å.

### Arg133 and Arg421 are essential for ER-exit, function and lipid-dependent stability

In addition to Arg287, Arg478 and Arg479 forming a lipid binding site at the dimer interface of UapA, MD simulations identified other, cytosolic-facing, residues that located on the outside of the core domain which have the potential to form interactions with annular lipids [20]. These residues, shown in Fig 5A, are Lys73 in the N-terminus just upstream of TMS1, Arg133, Tyr137 and Lys138 in loop L2, Lys212 in helix H1 of L4, and Arg421 in the border of L10 with TMS11. Of these residues, Arg421 is highly conserved in all NATs, whereas Lys73 and mostly Arg133 and Lys212 are highly conserved in fungal NATs. Tyr137 and Lys138 are not conserved in other NATs (S1 Fig).

**Fig 5.**
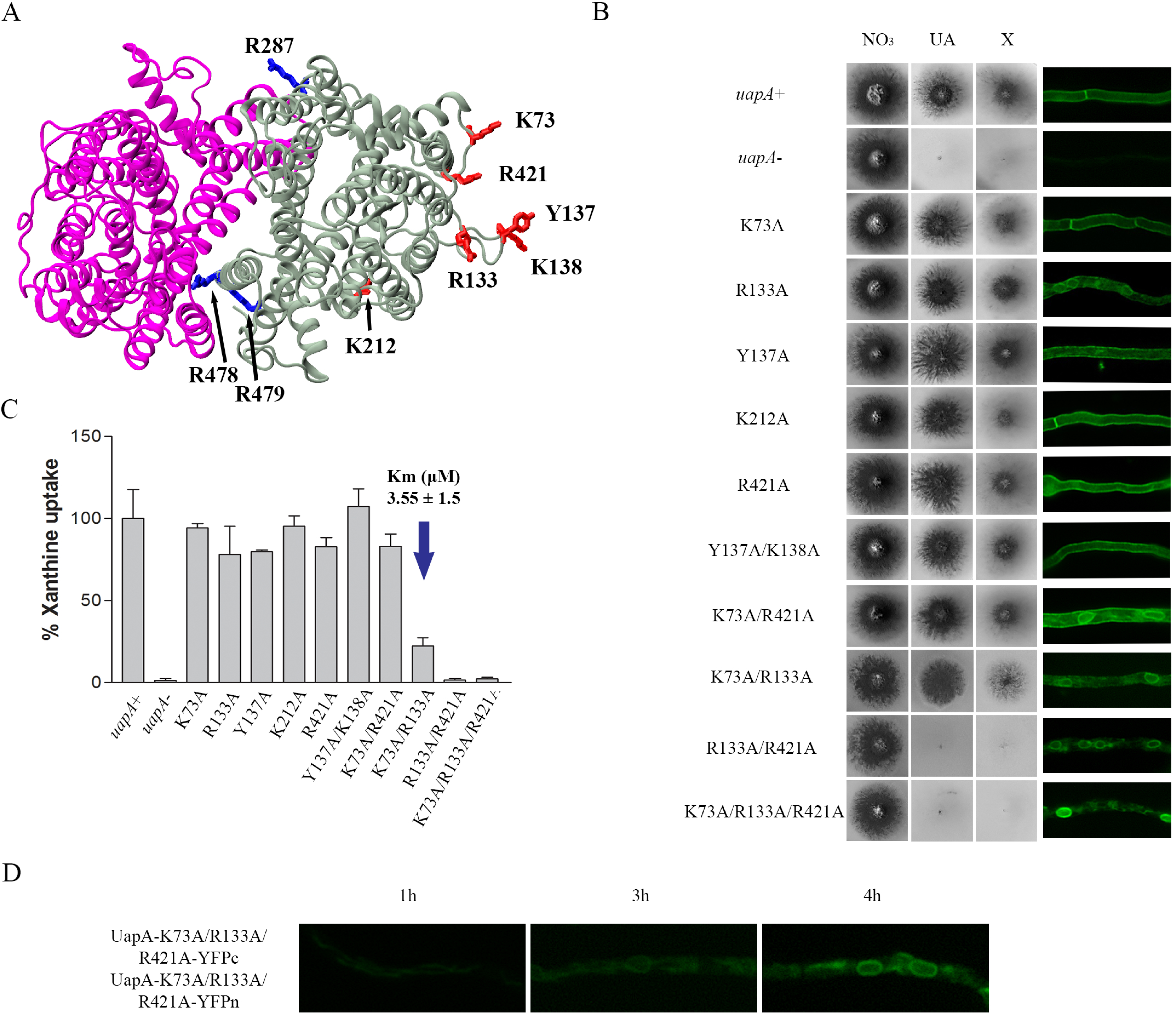
Arg133 and Arg421 are essential for ER-exit, function and lipid-dependent stability. (A) Topology of the predicted lipid binding sites near the dimer interface (blue) and those in the membrane-facing regions (red) of UapA dimer. (B) Growth tests of *A. nidulans* strains in minimal media supplemented with nitrate (NO3), uric acid (UA) or xanthine (X) as nitrogen source in 37°C (Left panel). Control strains and concentrations of supplements are as in Fig 2. All UapA mutant strains are isogenic to the negative and positive control strains, expressing *uapA* alleles tagged with *gfp.* Inverted fluorescence microscopy images shows localization of the GFP-tagged UapA constructs (Right panel). (C) Relative [^3^H]-xanthine transport rates of UapA mutants expressed as percentage of initial uptake rates (V) compared to the wild-type (*uapA^+^)* rate. [^3^H]xanthine uptakes were performed in 37°C. The *K*_m_ value (μM) for xanthine form mutant K73A/R133A is also indicated with the blue arrow. The results are averages of three measurements for each concentration point. SD was 20%. (D) Bimolecular complementation (BiFC) analysis of the K73A/R133A/R421A UapA mutant, performed as ppreviously described in Figs 1 and 3. Notice that the K73A/R133A/R421A retains the ability to reconstitute a flurosent signal in the ER, but not in the PM.

To investigate the potential functional role of the above residues we constructed mutants expressing all single, and selected double and triple Ala substitution. Fig 5B shows that the single mutations do not affect the growth on UapA substrates (xanthine or uric acid). Of the four double mutants constructed Y137A/K138A and K73A/R421A exhibited no UapA-related growth defect, while mutant K73A/R133A showed significantly reduced growth specifically on xanthine. Mutant R133A/R421A scored as an apparent total loss of function mutant, as it did not grow in either uric acid or xanthine. Finally, the triple K73A/R133A/R421A also scored as a total loss of function mutant. Overall, Arg133 and Arg421 proved very important for UapA function, while Lys73 when present in the context of R133A was critical for UapA specificity for xanthine, but not for uric acid. For each mutant version of UapA we also assessed localization to the PM, compared to the wild-type UapA, using the GFP epitope attached to UapA. This analysis (right panel in Fig 5B) showed that most single mutations and Y137A/R138A, which led to no defect in UapA transport activity, allowed normal localization of UapA in the PM, as probably expected. On the other hand, R133A and K73A/R421A mutants showed partial UapA retention in perinuclear ER membranes. Finally, mutants with apparently defective (K73A/R133A) or lost (R133A/R421A and K73A/R133A/R421A) transport activity showed partial or total retention into the ER. Growth tests and subcellular localization were in good agreement with measurements of rates of radiolabeled xanthine accumulation, which confirmed that the simultaneous presence of Arg133 and Arg421 is essential for transport activity, whereas Lys73 is critical for xanthine uptake only when Arg133 is also replaced by an Ala (Fig 5C). For the double mutant K73A/R133A which showed reduced growth on xanthine, we also measured the *K*_m_ for xanthine and showed that this was very close to that of the wild-type UapA (3.6 versus 5±2 μΜ), suggesting that reduced growth on xanthine is not assigned to reduced substrate binding.

We investigated whether the lack of UapA sorting out of the ER in the triple K73A/R133A/R421A mutant is related to its ability to dimerize. For this, we employed BiFC assays, as described before for the R287A/R478A/R479A mutant. Fig 5D shows that, rather surprisingly, the K73A/R133A/R421A mutant can apparently dimerize in the ER, as a strong fluorescence signal was reconstituted associated with ER membrane network. This was in line with western blots analysis that detected persisting dimeric forms of K73A/R133A/R421A (S2 Fig). These findings suggested that UapA dimerization is not sufficient for ER-exit and further sorting to the PM. This is also in agreement with the observation that lack or reduction of dimerization, seen in the R287A/R478A/R479A [20], did not interfere with proper ER-exit and sorting to the PM. Thus, functional dimerization and trafficking, processes seemingly affected by distinct lipid-interacting residues, are not necessarily related. Noticeably, attempts to purify the K73A/R133A/R421A protein showed that this version of UapA is prone to aggregation and is highly unstable (S3 Fig). Given that the K73A/R133A/R421A is very stable in total protein extracts (see S3 Fig), this means that instability occurs after removal of lipids during protein purification.

### Substitution T401P partially restores the lipid-dependent functional defects of K73A/R133A/R421A

We used a genetic approach to understand how residues Lys73, Arg133 and Arg421 might affect UapA sorting to the PM by isolating suppressor mutations that restored UapA-mediated growth on uric acid in the mutant K73A/R133A/R421A, as described earlier for the isolation suppressors of the R287A/R478A/R479A mutant. We obtained, purified and sequenced the *uapA* orf from 9 suppressors. Rather surprisingly, all proved to include the same single mutation, namely T401P, in addition to the original K73A/R133A/R421A triple mutation (Fig 6A). Noticeably, T401P was also isolated among the suppressors of the dimerization-defective R287A/R478A/R479A mutant. Growth tests showed that although T401P confers normal growth on xanthine and uric acid in the context of K73A/R133A/R421A (Fig 6B), this occurs by only partial restoration of UapA-mediated transport of these purines (Fig 6C). We also constructed by targeted mutagenesis plasmid vectors carrying K73A/R133A/R421A/T401P and T401P alone, introduced them in the *A. nidulans* strain lacking endogenous nucleobase transporters, and analyzed several purified transformants. Those carrying the quadruple mutation K73A/R133A/R421A/T401P behaved as the original suppressor, confirming that T401P is the causative mutation of suppressing the lack of function in R287A/R478A/R479A. Transformants expressing UapA-T401P behaved nearly as well as a wild-type UapA control, showing 70% transport rates and normal growth on xanthine or uric acid. Epifluorescence microscopy of K73A/R133A/R421A/T401P and T401P was in line with growth tests and uptake transport measurements. In particular, T401P in the genetic context of K73A/R133A/R421A partially restored sorting of UapA to the PM, while when present by itself it does not affect UapA localization to the PM (right panel in Fig 6B). High-copy transformants expressing K73A/R133A/R421A/T401P further confirmed that a major fraction of the mutant UapA translocates to the PM, despite some persistent retention in the ER (see Fig 6B). The positive effect of T401P in the context K73A/R133A/R421A was further shown by the isolation of fairly stable, detergent-solubilized, protein of K73A/R133A/R421A/T401P. Native mass spectrometry further showed that the purified K73A/R133A/R421A/T401P protein despite being mostly monomeric, it also formed a distinct population of dimers (S4 Fig). Thus, all evidence showed that T401P not only favors functional UapA dimerization in the R287A/R478A/R479A context, but also partially restores ER-exit, sorting and function in the K73A/R133A/R421A context.

**Fig 6.**
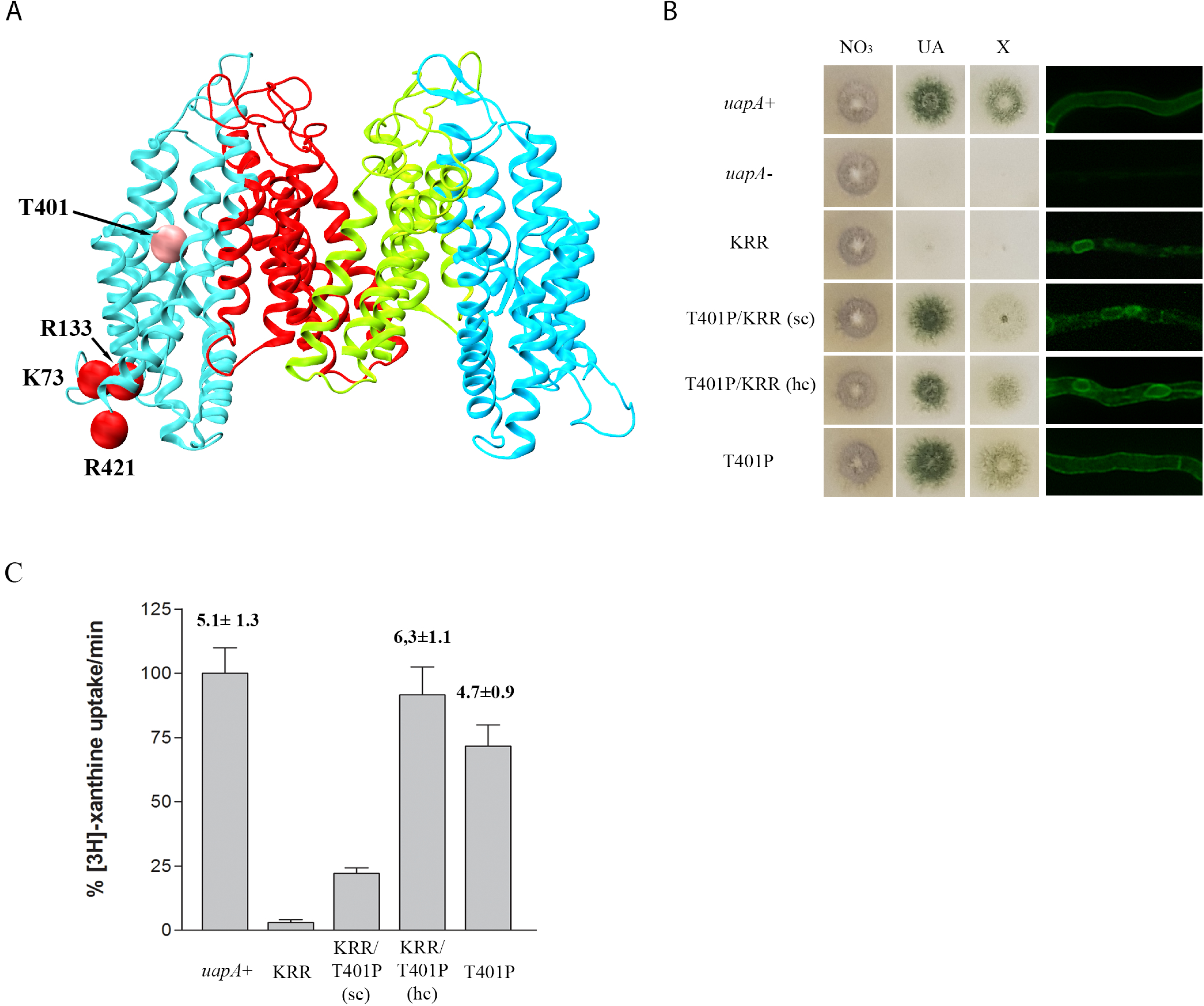
Substitution T401P partially restores the lipid-dependent functional defects of K73A/R133A/R421A. (A) Topology of amino acids modified in K73A/R133A/R421A suppressors. Core domains are colored light blue and dimerization domains red and green. Mutated amino acids in the original strain are shown with red spheres and in the suppressor with pink spheres. (B) Growth tests of K73A/R133A/R421A suppressors in minimal media supplemented with nitrate (NO3), uric acid (UA) and xanthine (X) as nitrogen sources in 37°C (Left panel. Control strains and supplement concentrations are as in previous Figs. All mutants are isogenic to the negative and positive control strains, expressing *uapA* alleles tagged with gfp. Inverted fluorescence microscopy images show localization of the GFP-tagged UapA constructs (Right panel). KRR depicts K73A/R133A/R421A. (C) Relative [^3^H]-xanthine transport rates of UapA mutant versions expressed as percentage of initial uptake rates (V) compared to the wild-type (*uapA+)* rate. [^3^H]-xanthine uptakes were performed in 37°C. *K*_m_ values (μM) for xanthine are shown at the top of histograms. Results are averages of three measurements for each concentration point. SD was 20%.

### Manipulation of a residue topologically equivalent to T401P leads to functional expression of a mammalian NAT homologue in *A. nidulans*

While *A. nidulans* or *Saccharomyces cerevisiae* have been successfully used to functionally express plant solute transporters (see early examples in [25–27]), including a UapA homologue [28], long-standing efforts of our group and many others have failed to functionally express metazoan solute transporters in model fungi. In all cases metazoan transporters are retained in the ER of fungi and often elicit UPR. The simplest explanation for this outcome is that the membrane environment of the fungal ER is incompatible with packaging metazoan transporters into COPII secretory vesicles [29]. Based on this idea and the fact that T401 proved to be a key residue in restoring lipid-dependent defects in UapA subcellular sorting and function, seemingly by increasing the compactness of the core domain of UapA, we thought we might achieve the functional expression of metazoan NATs in *A. nidulans* by manipulating similar key residues involved in interactions with lipids.

To test this idea, we used a characterized NAT homologue from rat, rSNBT1, which similarly to UapA acts as nucleobase transporter [22, 23]. Noticeably, UapA and rSNBT1 have different specificity and cation dependence, as rSNBT1 is a rather promiscuous Na^+^ symporter specific for pyrimidines (uracil, thymine) and most purines (hypoxanthine, guanine, xanthine and uric acid), while UapA is quite specific for xanthine and uric acid and utilizes H^+^ for co-transport. Previous attempts to express rSNBT1 or rSNBT1/UapA chimaeric transporters in *A. nidulans* or *S. cerevisiae* have failed, always due to total ER-retention (Anezia Kourkoulou, Christos Gournas, Sotiris Amillis, Bernadette Byrne and George Diallinas, unpublished). To putatively identify residue(s) in rSNBT1 that are topologically and functionally equivalent to T401 of UapA, we built a structural model by homology threading using the available UapA crystal structure (Fig 7A and S5 Fig). Candidate residues, equivalent to T401 in UapA, proved to be Asn390 and Gly391. Both are part of conserved short sequence motif Gly-Thr-Gly-**Asn**^390^-**Gly**^391^ present in all metazoan NAT members, irrespectively of their specificity [30]. In ascomycetes, the analogous sequence motif is Φ-Thr-Φ-**Thr-Pro** (Φ stands for aliphatic amino acid), while in basidiomyces and other more primitive fungi it is less well conserved, being Φ-Thr-Φ-Thr/Ser/Ala/Pro-Pro. In other words, what clearly distinguishes metazoa from fungi in this region is the replacement of the Asn residue (390 in rSNBT1) with Thr, and the last Gly (391 in rSNBT1) with a Pro.

**Fig 7.**
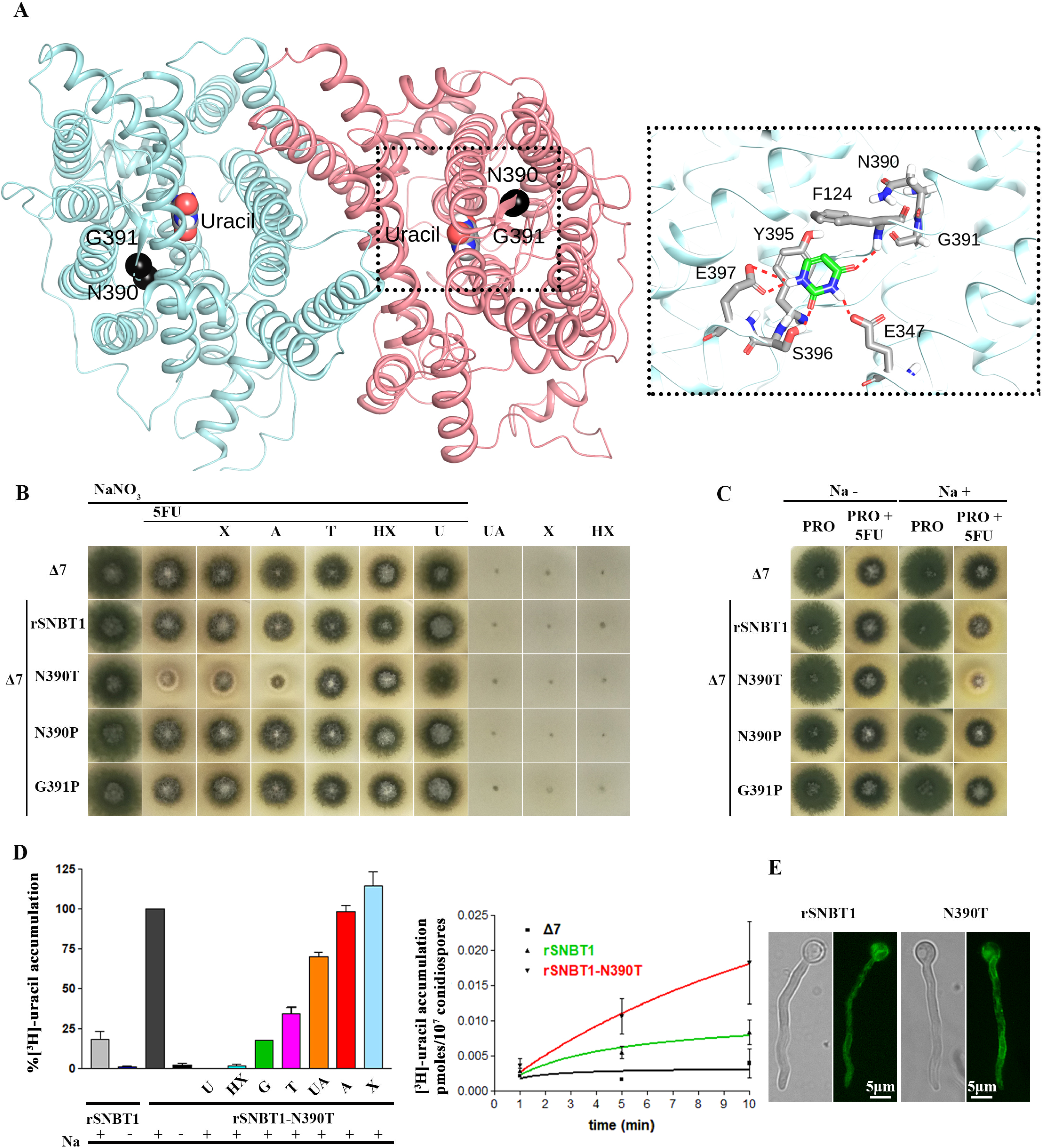
Manipulation of a residue topologically equivalent to T401P leads to functional expression of a mammalian NAT homologue in *A. nidulans*. (A) Homology modeling of the topology of rSNBT1, constructed using, as described in Materials and methods, the inward-facing conformation of the crystal structure of the UapA dimer. The two residues mutated and functionally analyzed, N390 and G391, are shown as black spheres. The location of uracil, the major substrate of rSNBT1, is also depicted, as determined by dynamic docking (left panel). In the right panel, a zoomed-out picture of the substrate binding site depicting the major interactions of uracil with specific residues. (B) Growth tests of isogenic *A. nidulans* strains expressing single-copy wild-type rSNBT1 or its mutated versions rSNBT1-N390T, rSNBT1-N390P and rSNBT1-G391P. A negative control strain (i.e. the recipient Δ7 strain that has null activity for nucleobase transport; see text) is included for comparison. Growth tests are performed at 37 ° C on MM supplemented with Na^+^ (100 mM NaCl). 10 mM NaNO_3_ is used as a control N source unrelated to purine transport activities in all tests scoring resistance/sensitivity to 5FU (rows 1-7). Rows 3-7 represent *in vivo* competion assays scoring the ability of excess purines (2mM) to compete with the uptake of 5FU (100 μΜ), and thus revert 5FU sensitivity. Χ is xanthine, A is adenine, T is Thymine, HX is hypoxanthine and U is uracil. Notice that T, HX and U competed 5FU uptake and suppressed sensitivity. Growth was also scored on MM containing UA (uric acid), X or HX, as sole nitrogen sources, none of which supported growth of the straisn tested (3 last rows). (C) Growth tests of *A. nidulans* of the same strains as in A, on MM plus proline as sole N source, supplemented or not with 100 mM NaCl. Notice that rSNBT1-N390T mediated 5FU sensitivity is dependent on the presence of Na^+^ supplementation. Notice also that in the presence of proline as N source, the wild-type rSNBT1 allele confers very moderate sensitivity to 5FU. (D) Left panel: [^3^H]-uracil (0.1 μΜ) accumulation in strains expressing *rSNBT1* and *rSNBT1-N390T,* performed in the presence or absence of 100mM Na^+^, and in the presence or absence of excess (2 mM) unlabeled nucleobases, after a period of 10 min incubation with radiolabeled substrate. [3H]-uracil accumulation in rSNBT1-N390T in the presence of 100mM Na^+^ and absence of unlabeled nucleobase is arbitrary taken as 100%. Right panel: relative [^3^H]-uracil (0.1 μΜ) transport accumulation in Δ7 (negative control), *rSNBT1* or *rSNBT1-N390T* strains as a time course. Uptake results are averages of three measurements for each concentration point. SD was 20%. (E) Inverted fluorescence microscopy images showing thje subcellualr localization of the GFP-tagged *rSNBT1* and *rSNBT1-N390T* constructs. Notice that the strains used for micoscopy are identical to those used in growth tests and uptake assays, as in all cases rSNBT1 sequences were tagged C-terminally with GFP (see Materials and methods).

Based on the above observations, we constructed and analyzed isogenic *A. nidulans* strains expressing the wild-type tSNBT1 or mutated versions with substitutions N390P, N390T or G391P in a genetic background lacking all endogenous transporters related to purine or pyrimidine uptake (see Materials and methods). Strains expressing wild-type or mutations N390P and G391P did not grow on purines and were resistant to 5-fluorouracil (i.e. a test for scoring uracil uptake), similarly to the recipient negative control strain lacking all endogenous nucleobase transporters. In contrast, the strain expressing rSNBT1-N390T showed clear sensitivity to 5-FU, and although it could not grow on any purine, 5-FU sensitivity could be competed in the presence of excess purines or pyrimidines that are known rSNBT1 substrates (e.g. hypoxanthine, uracil or thymine) (Fig 7B). Importantly, rSNBT1-dependent 5-FU sensitivity was Na^+^-dependent (Fig 7C), compatible with the physiological mechanism of functioning of rSNBT1 in rat [22, 23].

To further confirm that the phenotype observed in the relative transformants is due uniquely to the genetically introduced rSNBT1-N390T protein, we also analyzed the meiotic progeny of an rSNBT1-N390T transformant. *A. nidulans* undergoing meiosis during a process „selfing‟ [31] is prone to high recombination rates that often lead to loss of sequences introduced by transformation. S6 Fig shows that in an analysis of 28 meiotic progenital colonies of an original rSNBT1-N390T transformant, 21 colonies conserved the original sensitivity to 5-FU, while 7 colonies appeared as 5-FU resistant. Subsequent epifluorescence analysis of selected colonies showed that in all cases 5-FU sensitivity was conserved, a fluorescent signal from the rSNBT1-N390T protein tagged with GFP was also conserved. In contrast, all selected colonies that acquired resistance to 5-FU lost the fluorescent signal of the rSNBT1-N390T-GFP (not shown).

Finally, we also performed direct measurements of radiolabeled uracil accumulation or competition in the strain expressing rSNBT1-N390T, which further confirmed the functionality of the rat transporter in *A. nidulans* (Fig 7D). Noticeably, the low apparent transport capacity of rSNBT1-N390T in respect to some of is substrates (e.g. hypoxanthine or uric acid) in *A. nidulans* is very probably due by the observation that mutation N390T restores sorting of rSNBT1 to the PM only partially (Fig 7E).

## Discussion

It is becoming well-established that the physicochemical nature of lipid bilayers and specific lipid composition of membranes affect transporter folding, oligomerization, subcellular trafficking, function and turnover [8, 32–38]. For transporters conforming to the 5+5 inverted repeat (IR) or LeuT fold, similarities in structurally resolved lipid– protein interactions suggest common ways in which transporter structure and function are supported by lipid interactions [39]. These are likely to include stabilization of the inverted repeat topology, but also mechanistic roles as major determinants of the alternating access mechanism of secondary transporters. Noticeably however, the great majority of studies on transporter-lipid interactions have to date focused on prokaryotic transporters.

To our knowledge, our previous study on the UapA-lipid interactions still remains the only one focusing on a eukaryotic transporter [20]. In brief, we have shown that UapA, which primarily exists as a dimer, dissociates into monomers upon removal of tightly bound lipids, and that that dimer can be recovered by addition of PI or PE. Furthermore, as mutagenesis of tentative lipid-binding Arg residues 287, 478 and 479, predicted by MD, abolished lipid binding and function, we have proposed that PI/PE bind at specific sites in the dimer interface and thus stabilize the dimeric functional form of UapA [20]. The total lack of transport activity in R287A/R478A/R479A, despite the fact that in the mutant a degree of dimerization and normal sorting to the PM were still evident, suggested that lipid binding may also be directly essential for the mechanism of transport. Here, we studied further the role of Arg287, Arg478 and Arg479 and in parallel investigated the role of binding of annular lipids at distinct residues of UapA. We showed that Arg287, Arg478 and Arg479 are essential for early *de novo* formation the ER membrane, a process absolutely essential for transport activity, albeit not for sorting to the PM. In parallel, we identified distinct positively charged residues (Lys73, Arg133 and Arg421), exposed to the PM membrane bilayer, which are essential for ER-exit, sorting to the PM and transport activity, but apparently not essential for initial formation of dimers in the ER. Thus, the two sets of positively charged residues define two distinct sites of interaction of UapA with membrane lipids, both essential for function, albeit due to different reasons. The distinct defects caused by Ala substitutions at the dimer interface or those exposed to the inner side of the PM bilayer are well supported by epifluorescence microscopy, BiFC assays and native mass spectroscopy. Thus, while substitutions of Arg287, Arg478 and Arg479 did not affect UapA stability and sorting to the PM, substitutions of Lys73, Arg133 and Arg421 led to significantly protein instability and ER-retention. Interestingly, in no case, the mutant UapA versions studied elicited an Unfolded Protein Response (results not shown), suggesting that they do not lead to overall misfolding. This is particularly interesting mostly in the case of the K73A/R133A/R421A mutant, which is totally blocked within the ER membrane. This observation further suggests that interactions with annular lipids might be crucial for specific packaging into COPII secretory vesicles and ER-exit.

The most original finding of the present work stems from the isolation of genetic suppressors that partially restore defects caused due to modified interactions of UapA with either annular or specific lipids at the dimer interface. Most of the suppressors isolated mapped in the center of the core domain, and less frequently in the dimerization domain. No rational approach or MD studies could have predicted the functional importance of the residues identified via unbiased genetics. How these residues might correct defects in lipid binding became possible *a posteriori* with the help of MD and by taking into account the biophysical nature of residues introduced by suppressor mutations.

We can classify suppressors of the original dimerization mutant, R287A/R478A/R479A, into 3 types. Type I, which contains the majority of suppressors, map in the center of the core domain, in TMS2, TMS3, TMS4, TMS9, TMS1o and TMS11. All, except one, introduce residues with increased hydrophobicity and/or aromaticity (V150I, I157F, I157L, L192F, A396P, T401P, T401F or L431F). Only V153T introduces a polar residue, while S119T replaces a polar with longer residue of similar properties. MDs suggested that all these changes increase the strength of the relative TMS interactions and thus the compactness of the core domain and the stability of the protein. Type II includes L234M in TMS5 in the middle plane of the dimerization domain. This mutation could increase the strength of interactions between the dimerization and core domain. Type III includes E286Q and E286K at the end of the cytoplasmic-facing part of TMS6. The most logical scenario for these last two suppressors is that they replace directly the interactions with lipids of the mutated nearby Arg287 (i.e. in R287A). This is also in line with the fact that these are the strongest isolated suppressors in respect to UapA transport activity. Thus, our findings, especially those concerning type I and II suppressors, strongly suggest that by stabilizing the core, which is the motile part of the monomeric units that undergoes dynamic up and down elevator-like sliding, the dimer is also stabilized and thus function is restored.

Interestingly, all isolated suppressors of the „trafficking‟ mutant K73A/R133A/R421A proved to correspond to substitution T401P, a mutation that that also restored the dimerization mutant R287A/R478A/R479A. How this is achieved remains quite unclear, mostly because we still do not understand the molecular basis of the trafficking defect in the original mutant. Based on BiFC assays, western blot analysis and native mass spectroscopy, R287A/R478A/R479A was shown to form dimers, but these seem much more unstable and less abundant in respect to monomers, compared to wild type UapA [20]. Thus, despite the distinct defects caused by K73A/R133A/R421A and R287A/R478A/R479A triple, mutations in UapA subcellular localization, both seem to affect the stability and function of UapA via abolishment of essential but distinct interactions with specific or annular lipids. Apparently, mutation K73A/R133A/R421A was more critical than R287A/R478A/R479A for the packaging of UapA into COPII secretory vesicles and ER-exit [6], which might in turn suggest that annular lipid interactions are more important for the trafficking of UapA and other structurally similar eukaryotic transporters. In line with our results, it has been recently suggested that lipid binding around domain interfaces of the prokaryotic NhaA Na+/H^+^ exchanger are also involved in stabilizing the core domain during the conformational transitions required for transport by the elevator mechanism [38]. It has thus been speculated that elevator-type antiporters use a subset of annular lipids as structural support to facilitate large-scale conformational changes within the membrane. Other recent studies using native MS and functional assays have also demonstrated that protein-lipid interactions play a crucial role in stabilizing the dimer form of prokaryotic transporters conforming to the 7+7 inverted repeat topology found in transporter using the sliding elevator mechanism of transport [8, 17].

An impressive finding of our work was that a single mutation, T401P, proved to be a key residue in restoring defective interactions with either specific or annular lipids. We made use of this information and achieved, for the first time, the functional expression of a rat homologue of UapA, rSNBT1, in *A. nidulans*. The successful expression of rSNBT1 in *A. nidulans* strongly supports that the bottleneck in expressing metazoan transporters in fungi is proper folding in an environment of heterologous membrane lipid composition. Our achievement opens the way for further manipulations, via rational design or unbiased genetic screens, of lipid-binding residues in transporters for their functional expression and manipulation in genetically tractable model fungal systems, such as *A. nidulans* or *S. cerevisiae.* Additionally, the successful functional expressions of metazoan homologues in *A. nidulans* leads to new routes for identifying and studying the evolution of novel functions and substrate specificities in the NAT family, as for example, understand how human NAT homologues have evolved to become specific for vitamin C rather than nucleobases [30].

## Materials and Methods

### Media, strains and growth conditions

Standard complete (CM) and minimal media (MM) for *A. nidulans* growth were used. Media and supplemented auxotrophies were used at the concentrations given in http://www.fgsc.net. Glucose 1 % (w/v) was used as carbon source. 10 mM sodium nitrate (NO_3_^-^) or 10 mM proline was used as nitrogen source. Nucleobases and analogues were used at the following final concentrations: 5-fluorouracil (5-FU) 100 μΜ, uric acid (UA) 0.5 mM, xanthine (X) 1 mM, adenine (A) 2 mM, thymine (T) 2 mM, hypoxanthine (HX) 2 mM and uracil (U) 2 mM. All media and chemical reagents were obtained from Sigma-Aldrich (Life Science Chemilab SA, Hellas) or AppliChem (Bioline Scientific SA, Hellas). A *ΔazgA ΔuapA ΔuapC::AfpyrG pabaA1 argB2* mutant strain, named Δ3, was the recipient strain in transformations with plasmids carrying *uapA* alleles, based on complementation of the arginine auxotrophy *argB2.* A *ΔfurD::riboB ΔfurA::riboB ΔfcyB::argB ΔazgA ΔuapA ΔuapC::AfpyrG ΔcntA::riboB pabaA1 pantoB100* mutant strain, named Δ7, was the recipient strain in transformations with plasmids carrying the *rSNBT1* wild-type and mutated alleles, based on complementation of the pantothenic acid auxotrophy *pantoB100* [24]. *A. nidulans* protoplast isolation and transformation was performed as previously described [40]. Growth tests were performed at 25 or 37 ° C for 48 h, at pH 6.8.

### Standard molecular biology manipulations and plasmid construction

Genomic DNA extraction from *A. nidulans* was performed as described in FGSC (http://www.fgsc.net). Plasmids, prepared in *E. coli*, and DNA restriction or PCR fragments were purified from agarose 1% gels with the Nucleospin Plasmid Kit or Nucleospin ExtractII kit, according to the manufacturer‟s instructions (Macherey-Nagel, Lab Supplies Scientific SA, Hellas). Standard PCR reactions were performed using KAPATaq DNA polymerase (Kapa Biosystems). PCR products used for cloning, sequencing and re-introduction by transformation in *A. nidulans* were amplified by a high fidelity KAPA HiFi HotStart Ready Mix (Kapa Biosystems) polymerase. DNA sequences were determined by VBC-Genomics (Vienna, Austria). Site directed mutagenesis was carried out according to the instructions accompanying the Quik-Change® Site-Directed Mutagenesis Kit (Agilent Technologies, Stratagene). The principal vector used for UapA mutants was pAN510-GFP carrying a *gfp*-tagged *uapA* gene, as a template [41] and for *rSNBT1* mutants was a modified pGEM-T-easy vector carrying a version of the *gpdA* promoter, the *trpC* 39 termination region, and the *panB* selection marker [42]. For Bimolecular Fluorescence Complementation (BiFC) analyses, the N-terminal half of yellow fluorescent protein (YFPn; 154 amino acids of YFP), or the C-terminal half of YFP (YFPc; 86 amino acids of YFP) was amplified from plasmids PDV7 and PDV8 [43] and cloned into pAN510exp-*alcA_p_* or pAN520exp-*alcA_p_* [44] followed by cloning of the *uapA* ORF. UapA or *rSNBT1* mutations were constructed by oligonucleotide-directed mutagenesis or appropriate forward and reverse primers (S2 Table). Transformants arising from single copy integration events with intact UapA ORFs were identified by PCR analysis.

### Uptake assays

Kinetic analysis of UapA or rSNBT1 activity was measured by estimating uptake rates of [3H]-xanthine or [3H]-uracil uptake respectively (40-80 Ci mmol^−1^, Moravek Biochemicals, CA, USA), as previously described in [24]. In brief, [^3^H]-xanthine or [^3^H]-uracil uptake or competition by excess of other unlabeled substrates was assayed in *A. nidulans* conidiospores germinating for 4 h at 37° C, at 140 rpm, in liquid MM, pH 6.8. Initial velocities were measured on 10^7^ conidiospores/100 μL by incubation with concentrations of 0.2–2.0 μΜ of radiolabeled substrates at 37° C. The time of incubation was defined through time-course experiments and the period of time when each transporter showed linear increased activity was chosen respectively. All transport assays were carried out in triplets. Standard deviation was < 20%. Results were analyzed in GraphPad Prism software.

### Isolation and characterization of suppressor mutations

Suppressor mutations of 10^9^ conidiospores of strains R287A/R478A/R479A or K73A/R133A/R421A were obtained after 3 min 45 sec exposure at a standard distance of 20 cm from an Osram HNS30 UV-B/C lamp and subsequent selection of colonies capable of growing on MM containing uric acid as sole nitrogen source, at 25°C. Spores from positive colonies were collected after 6-8 days and further isolated on the same selective medium that was used to obtain the original colonies. Genomic DNA from 24 purified colonies was isolated and the *uapA* ORF was amplified and sequenced. In all cases the amplified fragments contained a new single missense mutation.

### Epifluorescence microscopy

Samples for standard epifluorescence microscopy were prepared as previously described [45, 46]. In brief, sterile 35 mm l-dishes with glass bottom (Ibidi, Germany) containing liquid minimal media supplemented with NaNO_3_ and 0.1% glucose were inoculated from a spore solution and incubated for 18 h at 25°C or for 8 h at 37 °C (rSNBT1). The samples were observed on an Axioplan Zeiss phase contrast epifluorescent microscope and the resulting images were acquired with a Zeiss-MRC5 digital camera using the AxioVs40 V4.40.0 software. Image processing and contrast adjustment were made using the ZEN 2012 software while further processing of the TIFF files was made using Adobe Photoshop CS3 software for brightness adjustment, rotation and alignment.

### Homology modeling of rSNBT1

The construction of a structural model of rSNBT1 was based on the crystal structure of the UapA in the inward-open conformation (PDB entry 5I6C). For this, we utilized the alignment shown in S5 Fig. The final model was built using PRIME software with an energy-based algorithm [47]. A loop refinement routine was also implemented.

### Induced Fit Docking of Uracil on rSNBT1

Protein preparation using OPLS2005 force field [48] and molecular docking was performed with the Schrödinger Suite 2018. After protein structure alignment with the crystal structure of UapA (PDB 5I6C) the binding pocket was defined by residues Phe124, Glu347, Tyr395, Ser396 and Glu397. Uracil was docked on the final structure from Homology Modeling, using the induced fit docking (IFD) protocol (Schrödinger Release 2018-1: Schrödinger Suite 2018-1 Induced Fit Docking protocol; Glide, Schrödinger, LLC, New York, NY, 2018; Prime, Schrödinger, LLC, New York, NY, 2018.), which is intended to circumvent the inflexible binding site and accounts for the side chain and backbone movements upon ligand binding [49].

### Molecular Dynamics

UapA (wild-type or mutated when discussed) or rSNBT1 homologue dimers were inserted into a lipid bilayer using the CHARMM-GUI tool [50]. The resulting system was explicitly solvated using the TIP3P water model [51] and neutralized by the addition of Na^+^ and Cl^−^ counter ions at concentration of 0.15 M. The lipid bilayer utilized was composed of 20% ergosterol, 9% POPC, 12% DYPC, 9% YOPC, 6% POPE, 3% DYPE, 5% YOPE, 3% DOPE, 19% POPI and 14% PYPI, as described previously [20]. All UapA mutations were constructed by utilizing the CHARMM-GUI’s initial step “PDB Manipulation Options”. The N-terminal residues were always methylated and the C-terminus residues were always amidated. Molecular dynamic (MD) simulations were performed with GROMACS 2018 [52] using the all-atom force field CHARMM36 [53]. Periodic boundary conditions were used. Long-range electrostatic interactions were treated with Particle Mesh Ewald (PME) method. Non-bonded interactions were described with a Lennard-Jones potential with a cut-off distance of 1 nm and an integration step of 2 fs was implemented. The system was progressively minimized and equilibrated using the GROMACS input scripts generated by CHARMM-GUI and the temperature and pressure was held at 303.15 K and 1 bar respectively [54]. The resulting equilibrated structures were then used as an initial condition for the production runs of 100 ns with all the constraints turned off. Production runs were subsequently analyzed using GROMACS tools and all images and videos were prepared using VMD software [55].

### *In silico* mutation of I157F, L192F and L431F on UapA

Staring from the crystal structure of UapA manual mutation of I157F, L192F and L431F was performed using “mutation” command on Maestro v11.5 (Schrödinger Release 2018-1). Each resulting structure was inserted to Protein Preparation Wizard Workflow as implemented on Maestro v11.5. Restrained minimization was converged when heavy atoms RMSD was greater than 1 Å.

### UapA K73A/R133A/R421A (T401P) Expression and Purification

Expression and purification of the UapA K73A/R133A/R421A and K73A/R133A/R421A/T401P mutants was carried out as described previously [20]. In brief the mutants were individually expressed as C-terminally GFP8His tagged constructs in *S. cerevisiae* FGY217 cells (12L), using vector pDDGFP2. Yeast cells were incubated at 30 °C with shaking at 300 rpm to an OD_600_ of 0.6. Galactose was then added to the cultures to a final concentration of 2 % to induce UapA expression. After incubation for a further 22 hours, the cells were harvested by centrifugation and resuspended. Cells were broken in a Constant Systems cell disruptor and the membranes isolated by centrifugation. The membranes were resuspended, flash-frozen, and stored at −80 °C. Membranes were solubilized for 1 hour in n-dodecyl-β-D-maltoside (DDM). Unsolubilized membranes were removed by centrifugation. The supernatant was incubated with Ni^2+^-NTA resin for 2 hours. The His-tagged UapA bound to the resin was then washed with buffers containing 10 mM and 30 mM imidazole to remove contaminants. UapA was eluted with buffer containing 250 mM imidazole before overnight dialysis to dilute the imidazole from the protein sample. During dialysis, the protein was cleaved using a His-tagged TEV protease. The sample was run through a His-trap column, from which UapA was eluted in the flow through, in order to remove the His tagged GFP and TEV. The sample was then loaded onto a size exclusion chromatography column. Fractions containing monodisperse UapA were analyzed by SDS-PAGE and concentrated to 20 μM, flash-frozen, and stored at −80 °C.

### Native Mass Spectrometry (MS) of UapA

Native MS of UapA K73A/R133A/R421A/T401P was carried out as described previously [20]. In brief, UapA was buffer exchanged into MS buffer (250 mM EDDA (pH 6.3), 0.014% DDMLA (v/v), 10 mM L-serine) to a UapA concentration of 20 μM using Micro Bio-Spin 6 columns (Bio-Rad). UapA was loaded into gold coated capillaries and the protein sprayed into a Synapt G2-Si (Waters) by nano-electrospray ionization. The following conditions were used in the mass spectrometer for optimal peak resolution: capillary voltage +1.3-1.5 kV, sampling cone voltage 150 V, trap collision energy (CE) 200 V, transfer CE 0 V, backing pressure 3.88 mbar, trap and transfer pressure (argon) 1.72e-2 mbar, ion mobility cell pressure (nitrogen) 2.58 mbar. The mass spectrometer was calibrated using cesium iodide. Spectra were recorded and processed using Masslynx 4.1 software (Waters). The relative abundances of each oligomeric state were quantified by UniDec [56] as described previously [20].

## Acknowledgments

This work was supported by a “Stavros S. Niarchos Foundation” grant to A.K and G.D, and by computational time granted from the Greek Research & Technology Network (GRNET) in the National HPC facility -ARIS-under project NCS1_Mechanism (pr006040). This work was also supported by a Biotechnology and Biosciences Research Council grant, BB/N016467/1. E.P. is the recipient of an Imperial College London Institute of Chemical Biology Engineering and Physical Sc

## Author Contributions

Conceptualization: G.D, B.B

Formal analysis: G.D, B.B, E.M.

Funding acquisition: G.D, B.B

Investigation:-Methodology: A.K., P.G., G.L., M.D., E.P.

Project administration: G.D, B.B

Resources: G.D, B.B, A.P, E.M.

Supervision: G.D, B.B, A.P., E.M.

Writing-original draft: G.D, E.M.

Writing – review & editing: G.D, B.B

## Supporting information

**S1 Fig.**
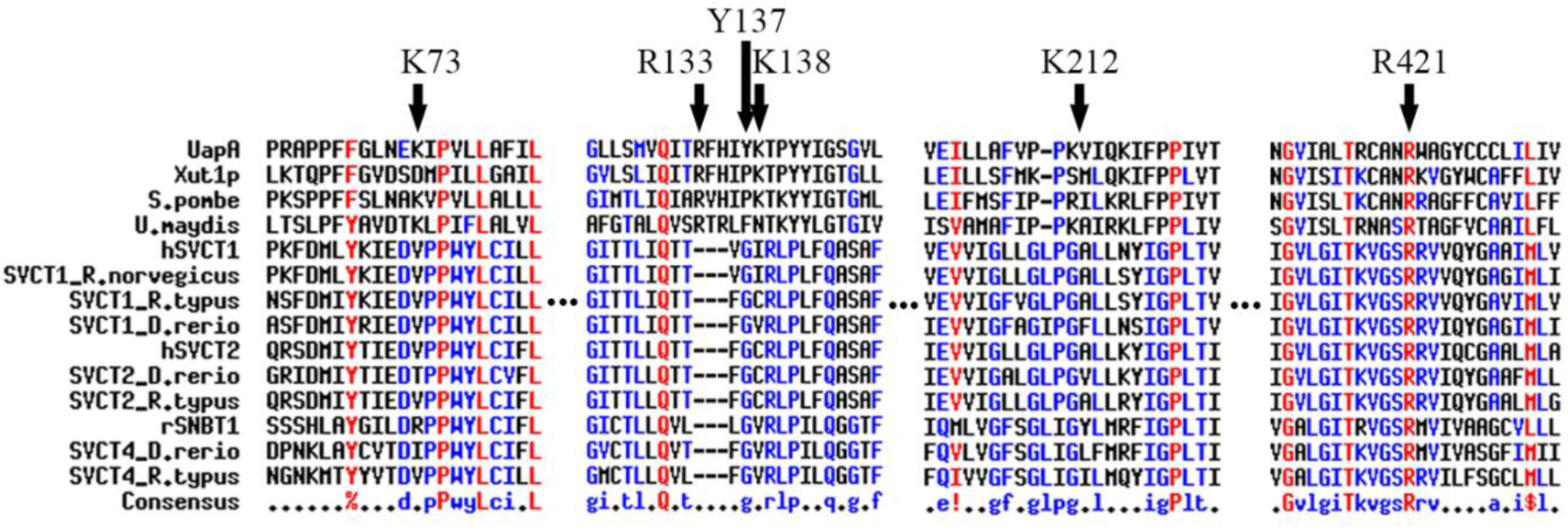
Multiple sequence alignments of specific segments of UapA and selected homologues from fungi and metazoa depicting the conservation of residues K73, R133, R137, K138, K212 and R421 studied in the recent work. Xut1 is a *Candida albicans* orthologue. hSVCT1, hSVCT2 and rSNBT1 are discussed in detail in the text (human L-ascorbate and rat nucleobase transporters respectively). Full species names and specific protein codes are as follows: *Schizosaccharomyces pombe* (NP_593513.1), *Ustilago maydis* (XP_011390165.1)*, Rattus norvegicus* (XP_006254664.1 and NP_059012.2), *Danio rerio* (NP_001166970.1, XP_005169157.1 and NP_001013353.1), *Rhincodon typus* (XP_020385431.1 and XP_020391140.1)

**S2 Fig.**
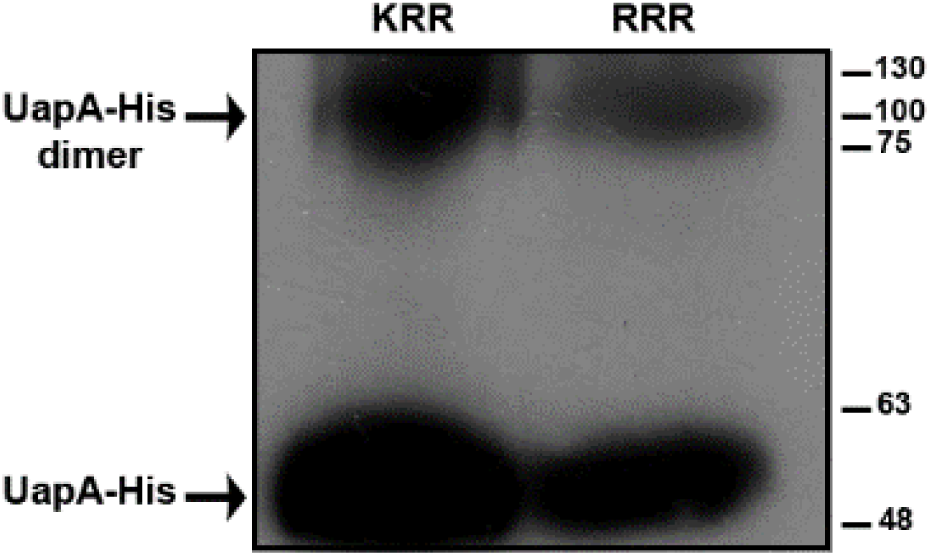
A significant fraction of R287A/R478A/R479A and K73A/R133A/R421A mutants forms tight UapA dimer. Western blot analysis, using anti-His antibody, of total protein extracts from strain expressing R287A/R478A/R479A-His (RRR) or K73A/R133A/R421A-His (KRR) expressed under the control of the regulatable promoter *alcA*p. The two strains were incubated for 20 h under derepressed (0.1% fructose as C source) and induced (0.1% ethanol) conditions for 4h before protein extraction. For SDS-PAGE [10% (w/v) polyacrylamide gel] proteins were loaded directly to the gel with 3x sample loading buffer without SDS [glycerol, 50x running buffer (Tris Base, Glycine), bromophenol blue]. Western blot immunodetection of the UapA mutants, showed a prominent band at the position corresponding to monomeric UapA-His (∼55kDa) and a high molecular weight band (∼110kDa), corresponding to the expected size of dimeric form, similar to the one usually detected for wild-type UapA.

**S3 Fig.**
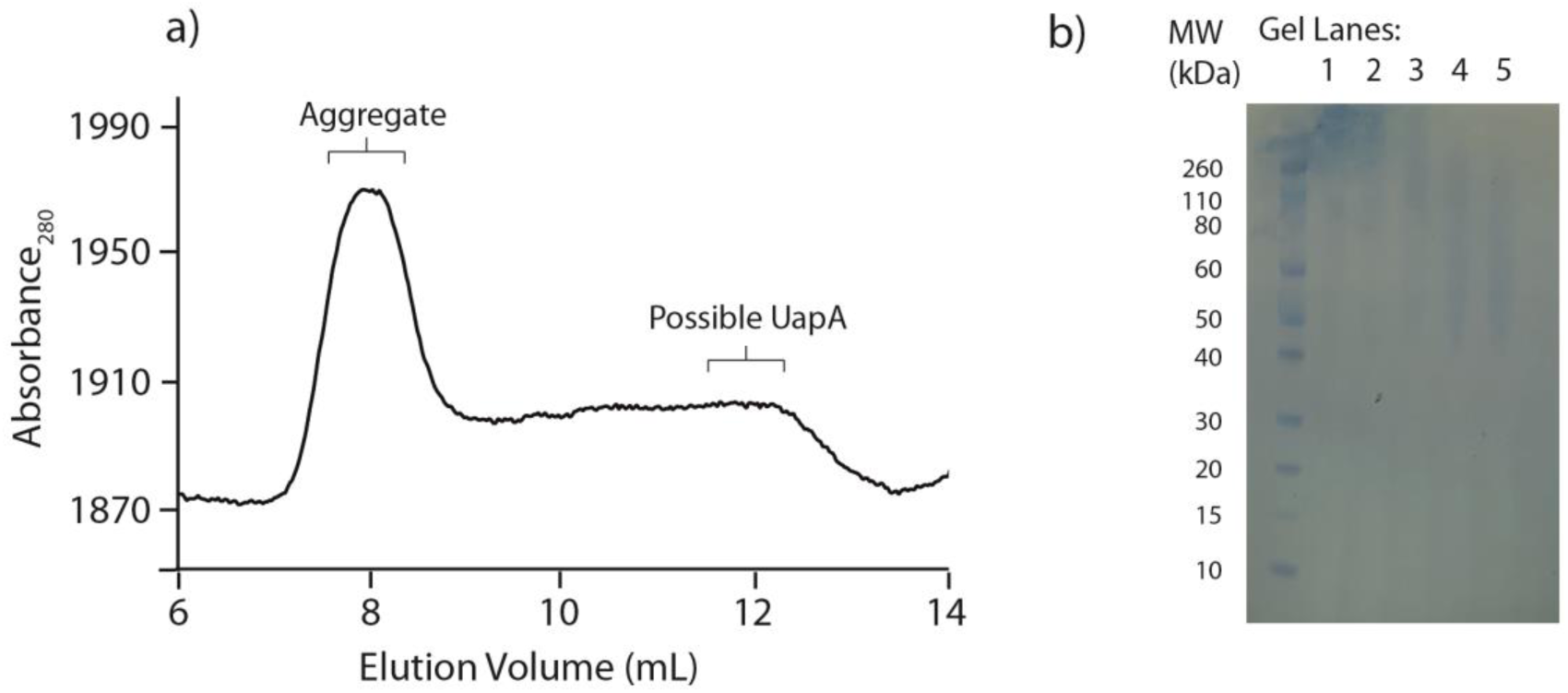
Purification of UapA K73A/R133A/R421A (a) SEC profile of the mutant. Fractions used for SDS-PAGE gel lanes are indicated with labels. (b) SDS-PAGE gel from SEC. Gel lanes 1-2: Aggregate; 3-5: Possible UapA. Note the smear at the top of lanes 1 and 2 due to aggregated protein.

**S4 Fig.**
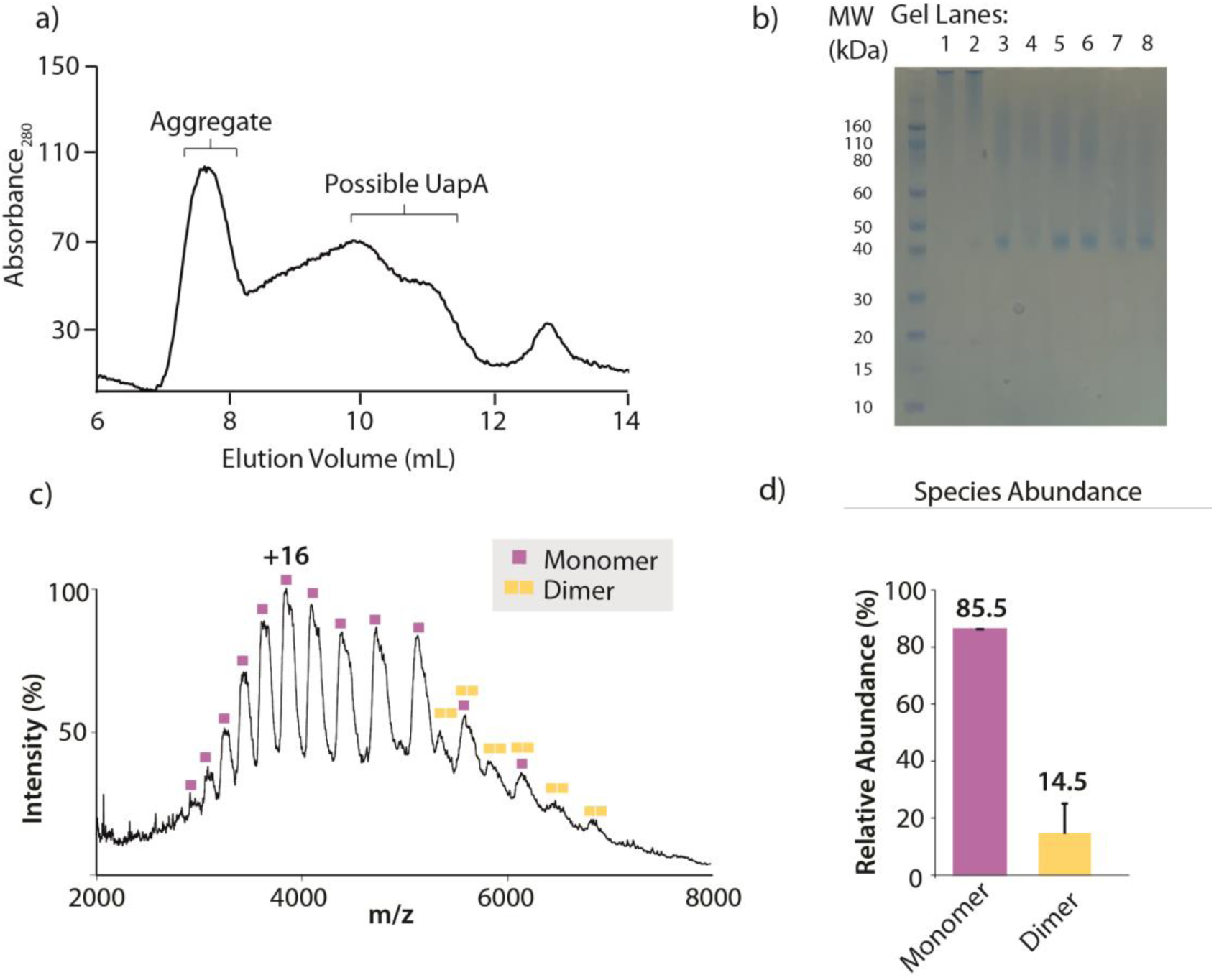
Purification and native MS of UapA K73A/R133A/R421A/T401P. (a) SEC profile of the mutant. Fractions used for SDS-PAGE gel lanes are indicated with labels. (b) SDS-PAGE gel from SEC. Gel lanes 1-2: Aggregate; 3-8: Possible UapA. The peak at 13 mL is due to DDM eluting from the column. (c) Native mass spectrum of UapA K73A/R133A/R421A/T401P with monomer and dimer peaks indicated. (d) Relative abundance of monomer and dimer was calculated using UniDec (Marty et al, 2015). Average abundance was calculated from three repeats carried out under identical conditions in the mass spectrometer.

**S5 Fig.**
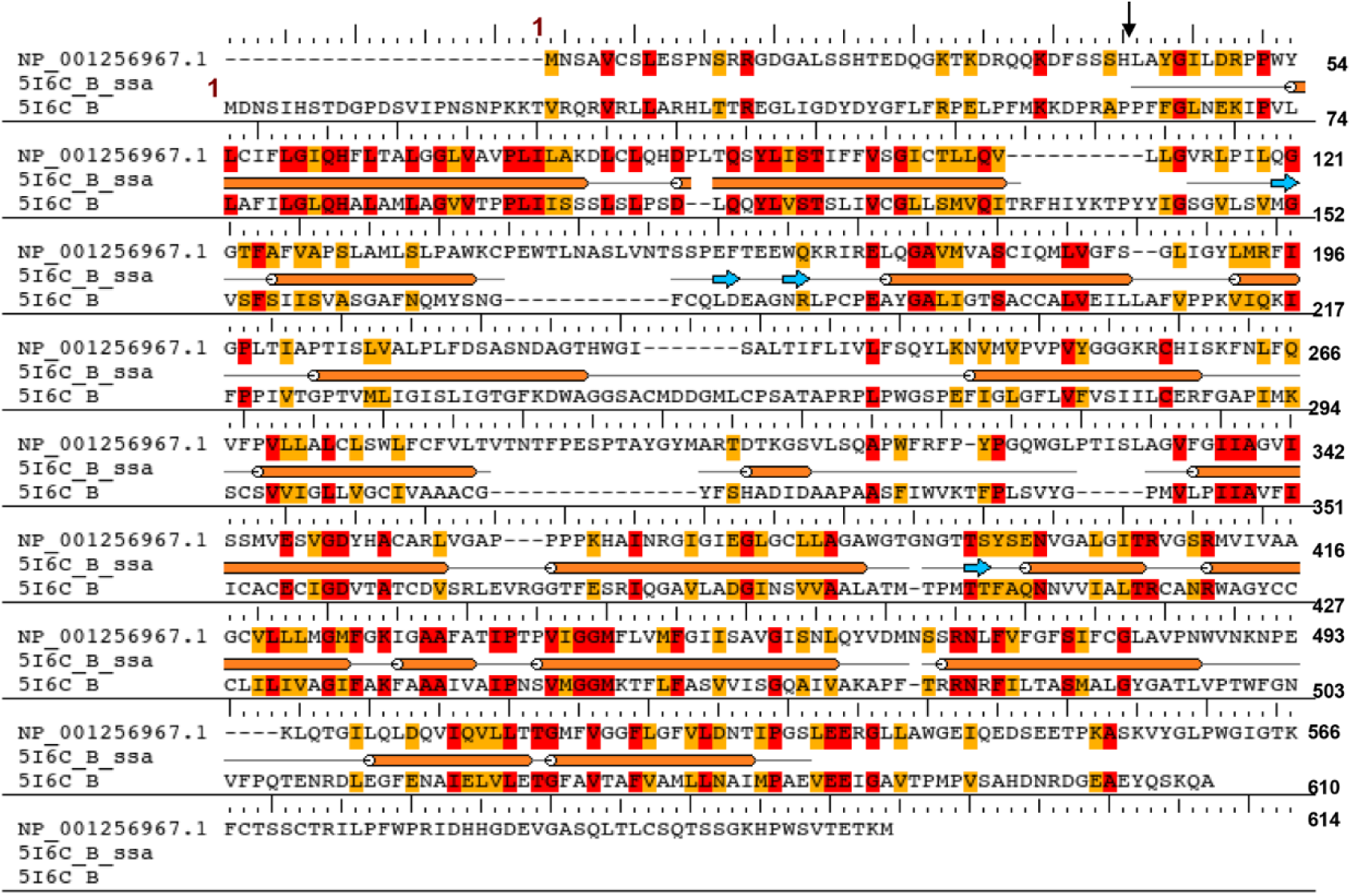
Alignment of rSNBT1 vs UapA utilized for the Homology Modeling of rSNBT1. Number 1 indicates the starting amino acid in the sequence. The arrow indicates the starting amino acid of UapA crystal structure.

**S6 Fig.**
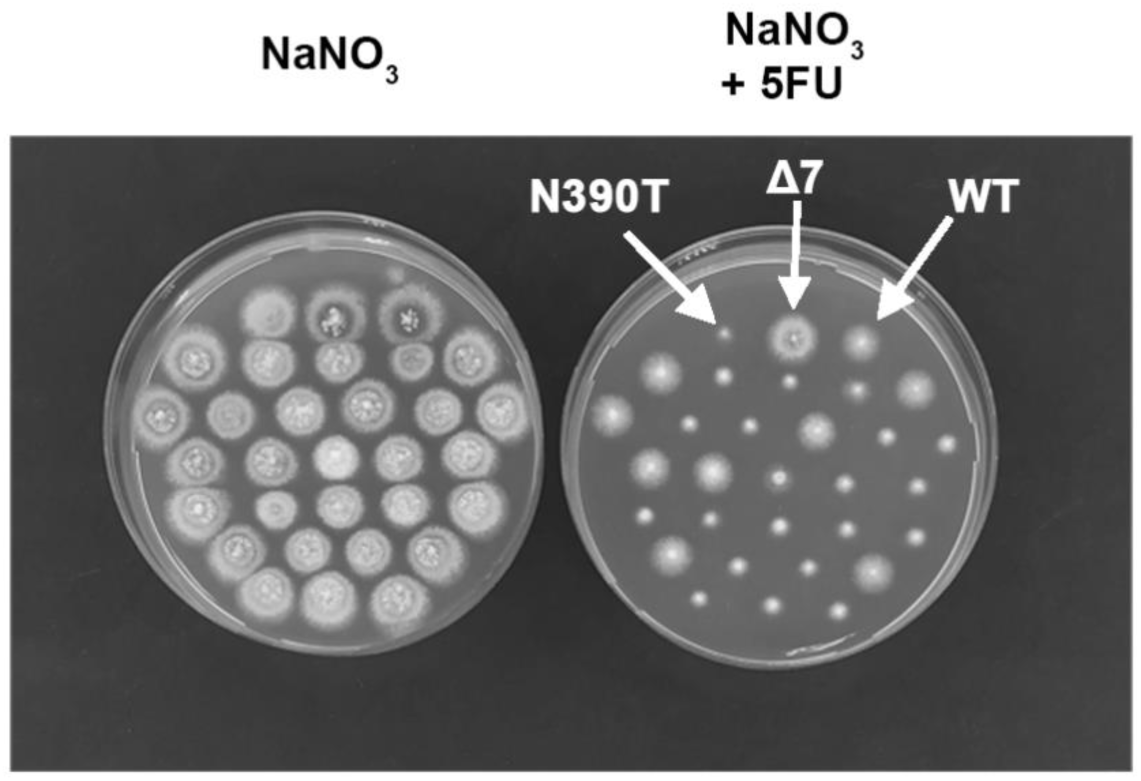
Co-segregation of 5-FU sensitivity with genetic loss of rSNBT1-N390T activity. The figure shows a growth test on minimal media, supplemented with 100 NaNO3 mM or 100 mM NaNO3 plus 100 μM 5-fluorouracil (5-FU) of 29 meiotic progeny colonies originating from individual ascospores from a ‘selfed’ cleistothecium of a strain that contained a genome-integrated single plasmid copy of rSNBT1-N390T. Isogenic control strains are a strain with total deletions in all major purine transporters (Δ7 negative control), a Δ7 transformant expressing rSNBT1-N390T-gfp and a standard wild-type strain expressing all known purine transporters. Notice that 8 colonies are resistant to 5-FU, contrasting the sensitivity of the parental strain. These colonies were subsequently shown to have lost the rSNBT1-N390T sequences, apparently due to genetic excision of integrated plasmid sequences, a frequent meiotic phenomenon in *A. nidulans* [31].

**S1 Table.** Profile and frequency of isolation of suppressors R287A/R478A/R479A.

| **Mutation** | **Location** | **Domain** | **Number of isolates** | **Codon change** | **Number of changed nucleotides** |
| --- | --- | --- | --- | --- | --- |
| **S119T** | TMS2 | Core | 1 | TCG→ACG | 1 |
| **V150I** | TMS3 | Core | 9 | GTT→ATT | 1 |
| **V153T** | TMS3 | Core | 1 | GTC→ACC | 2 |
| **I157F** | TMS3 | Core | 3 | ATC→TTC | 1 |
| **I157L** | TMS3 | Core | 3 | ATC→CTC | 1 |
| **L192F** | TMS4 | Core | 2 | TTG→TTT or TTC | 1 |
| **L234M** | TMS5 | Dimerization | 2 | CTG→ATG | 1 |
| **E286K** | TMS6 | Dimerization | 6 | GAA→AAA | 1 |
| **E286Q** | TMS6 | Dimerization | 2 | GAA→CAA | 1 |
| **A396P** | TMS9 | Core | 1 | GCC→CCC | 1 |
| **T401F** | TMS10 | Core | 1 | ACC→TTC | 2 |
| **T401P** | TMS10 | Core | 1 | ACC→CCC | 1 |
| **L431F** | TMS11 | Core | 6 | TTG→TTT or TTC | 1 |

**S2 Table.**
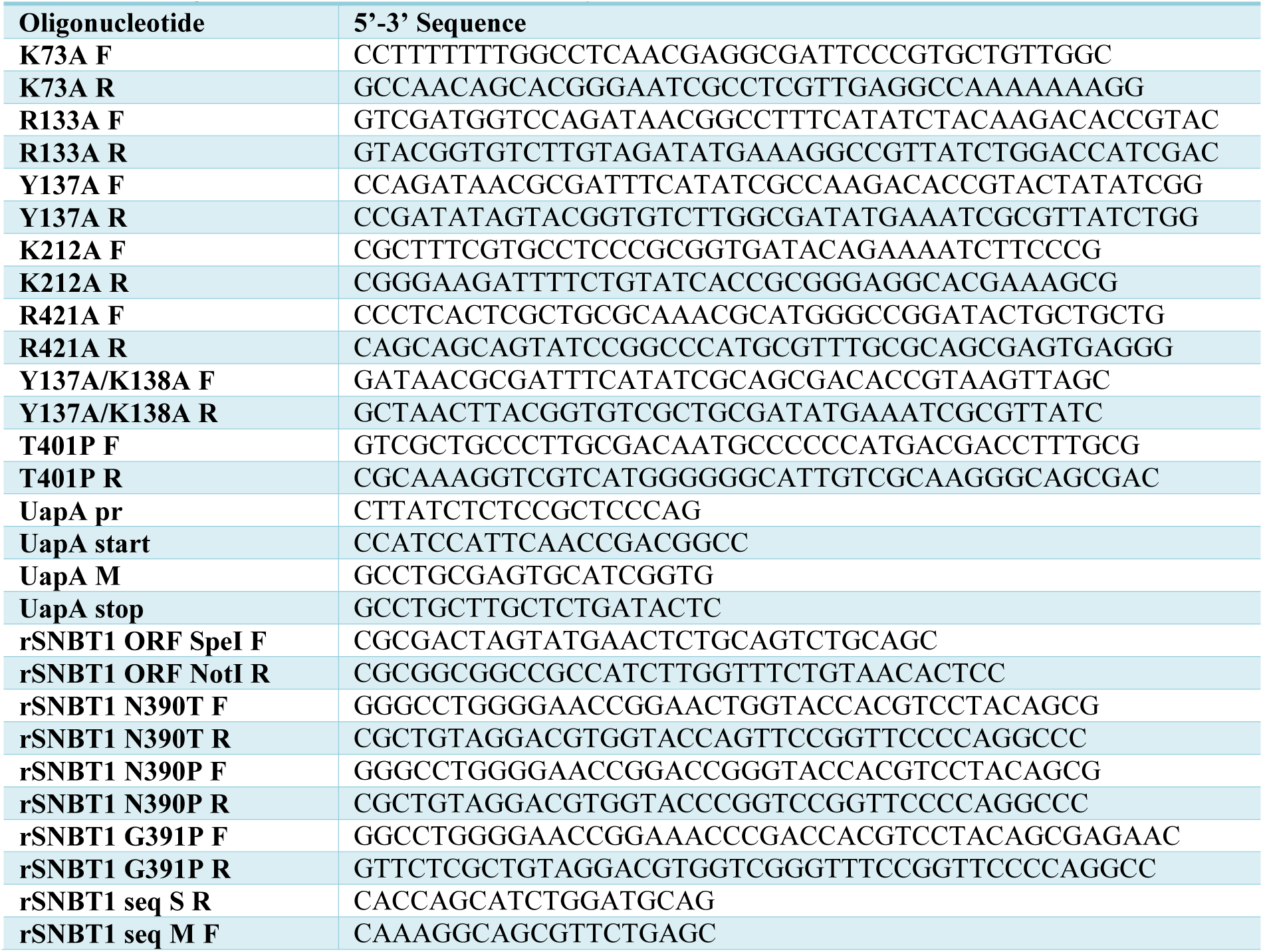
Oligonucleotides used in this study.

